# Rational arbitration between statistics and rules in human sequence processing

**DOI:** 10.1101/2020.02.06.937706

**Authors:** Maxime Maheu, Florent Meyniel, Stanislas Dehaene

## Abstract

Detecting and learning temporal regularities is essential to accurately predict the future. A long-standing debate in cognitive science concerns the existence of a dissociation, in humans, between two systems, one for handling statistical regularities governing the probabilities of individual items and their transitions, and another for handling deterministic rules. Here, to address this issue, we used finger tracking to continuously monitor the online build-up of evidence, confidence, false alarms and changes-of-mind during sequence processing. All these aspects of behaviour conformed tightly to a hierarchical Bayesian inference model with distinct hypothesis spaces for statistics and rules, yet linked by a single probabilistic currency. Alternative models based either on a single statistical mechanism or on two non-commensurable systems were rejected. Our results indicate that a hierarchical Bayesian inference mechanism, capable of operating over distinct hypothesis spaces for statistics and rules, underlies the human capability for sequence processing.

## Main text

From weather to traffic lights, many real-life processes unfold across time, forming sequences of events characterised by some inner structure. The ability to learn such sequential regularities is essential to agents navigating real-life environments because it enables them to make predictions about the future ^1–3^. Past research indicates that the brain constantly entertains such predictions ^4–6^ which it leverages to promote a more efficient processing of incoming information as well as to improve decision-making and behavioural control ^7–9^.

In order to make accurate predictions, human observers must solve two difficult problems. First, they should *detect when a regularity appears* ^10–14^. Second, because various types of regularities exist, observers must also *identify the kind of process generating the regularity*. In theory, the number of hidden generative processes is infinite. Here, we study a simple distinction between statistical biases and deterministic rules, which relates to the strength of predictions they afford (*Fig. 1a*). Statistical biases allow uncertain, yet better-than-chance, predictions. For instance, looming dark clouds are generally followed by rain, but sun may also appear on some rare occasions. Deterministic rules, on the other hand, allow sure predictions. For instance, at a French traffic light, only the orange light (warning signal) can follow the green one (go signal).

**Fig. 1.**
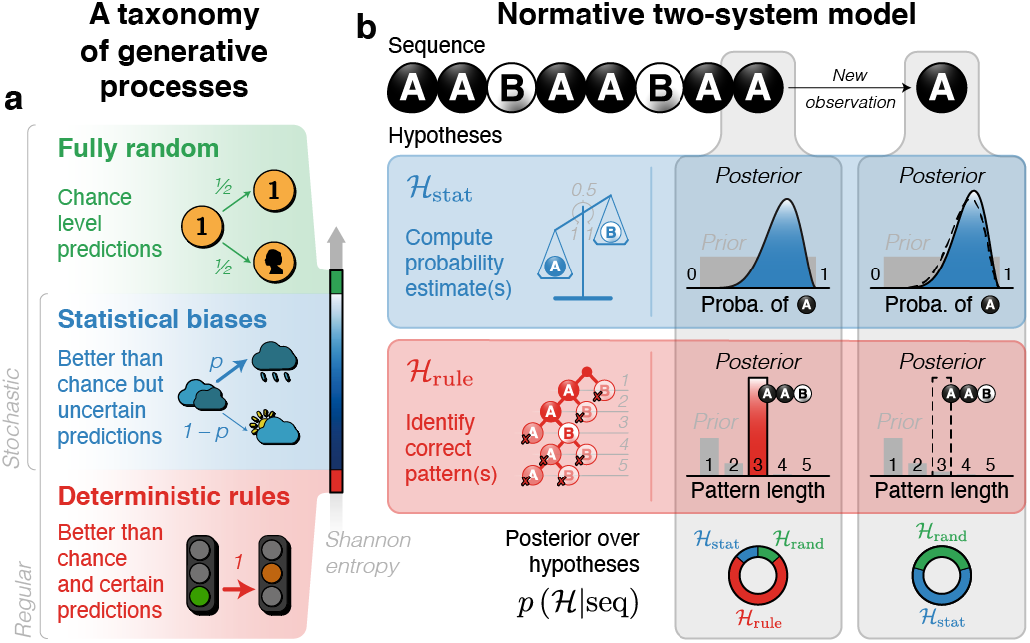
A taxonomy of regularities and a normative model of how those regularities can be inferred from a sequence of observations. **a**, *A taxonomy of regularities*. A first distinction separates random (i.e. unpredictable) from non-random (i.e. predictable) processes. A second distinction, within non-random processes, separates statistical biases (i.e. affording only uncertain predictions) from deterministic rules (i.e. affording certain predictions). Shannon entropy quantifies the unpredictability of a process and therefore differs between those different types of processes. **b**, *Normative two-system model*. An example sequence of binary observations is presented to the model which arbitrates between different hypotheses by means of probabilistic inference. For simplicity, we supposed here that there is no volatility: the full binary sequence corresponds to a single generative process, which can be either random, or exhibit a statistical bias (here, in the frequency of items) or follow a deterministic rule (here, the repetition of a fixed pattern of at most five items). Following the first 8 observations, the statistical bias hypothesis estimates than As are overall more likely than Bs. By contrast, the deterministic rule hypothesis estimates that the sequence can be described as the repetition of the AAB pattern. Computing the posterior probability of each hypothesis reveals that, at this point, the deterministic rule hypothesis is the most likely hypothesis. Note, however, that a single new observation (here, the final A) suffices to discard the deterministic rule hypothesis.

In our taxonomy, deterministic rules are an extreme case of statistical biases, in which the probability of the next event is 0 or 1 (*Fig. 1b*). However, here, we test the hypothesis that humans treat statistics and rules as fundamentally different, corresponding to two distinct hypothesis spaces. Indeed, those two types of regularity allow predictions of a different nature (uncertain vs. certain), they enable computations of different kinds (which we detail later). The statistics versus rules debate has long divided cognitive scientists ^15–19^, and as a result, they have been explored in mostly distinct lines of research. On the one hand, detecting, estimating or leveraging statistical biases is at the heart of studies on probability learning ^14,20–26^, reinforcement learning ^27–33^, and artificial grammar learning paradigms involving Markov processes ^34–37^. On the other hand, deterministic rules were embedded, for instance, in paradigms in which repeated patterns have to either be detected ^10,13,38–41^ or reproduced ^42–45^. Deterministic rules more complex than simple repetitions have also been explored ^2,38,46^. Studies that combine or compare both kinds of regularities appear as exceptions ^47–53^.

In this article, we propose a new paradigm in order to jointly study the problems of detecting when a regularity emerges, and of identifying its type. This approach offers opportunities to bridge gaps between previous studies which investigated those aspects one at a time. Notably, this approach enables to test a new theory, which assumes that the detection and identification of regularities should rely on a hierarchical Bayesian inference over a minimum of three distinct hypothesis spaces: fully random, statistical bias, and deterministic rule. This new proposal makes several predictions, not only in terms of subjects’ performance in detecting regularities and discriminating between types of regularities, but also regarding the strength and fluctuations of their beliefs, the specific dynamics associated with the detection, and even the type of errors people make. We designed a new experiment to test those predictions, and compared the results with alternative models of regularity detection.

## Results

### Experimental design

To test our theoretical proposal, we designed a novel behavioural experiment in which we presented human subjects with binary auditory sequences (200 observations, for 1 minute each; *Fig. 2a*; *Extended Data Fig. 1* for all sequences). All sequences started with a random process (i.e. like tossing a fair coin). In about two thirds of the cases, after a delay of variable length, we introduced either a statistical bias (in the first-order transition probabilities) or a deterministic rule (the repetition of a particular pattern of length-4 to −10). In the remaining cases, the sequences remained fully random until the end. Subjects were fully instructed about this task structure and were asked to slide their finger towards one of three locations, corresponding to the generative process underlying a given sequence (*Fig. 2b,c*). In order to monitor the dynamics of inference, we used a continuous finger tracking system. Subjects also provided a detailed subjective report after each sequence. Our design therefore combines two aspects: detecting when a regularity emerges and identifying its type (i.e. statistic or rule).

**Fig. 2.**
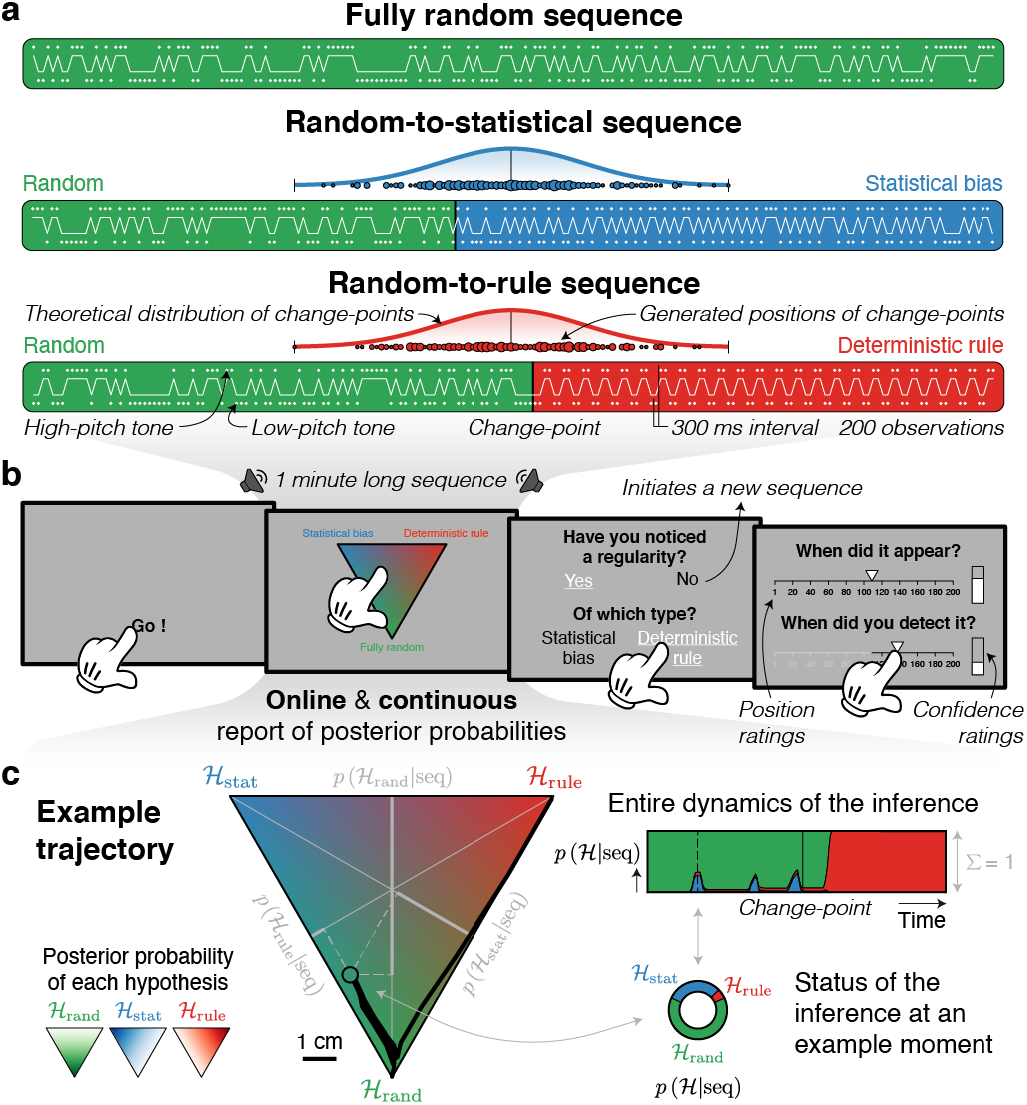
Behavioural task. **a**, *Example sequences*. Sequences were of three different types: random from the beginning to the end or composed of an initial random part and a second non-random part that could either be produced by a statistical bias (here, a bias toward alternation) or a deterministic rule (here, the repetition of the AABB pattern). These sequences were composed of 200 low- and high-pitch tones (interval of 300 ms) that were randomly assigned to observations A and B at the beginning of each sequence. The position of the change-point was drawn from a gaussian distribution centred on the 100^th^ observation, with a standard deviation of 15 observations, and truncated from the 55^th^ and 145^th^ observations; generated change-point positions are shown as small circles whose size indicates their frequency. **b**, *Timecourse of a trial*. Each trial was self-initiated. During sequence presentation, subjects were asked to move their finger within a triangular arena whose vertices correspond to the three possible generative processes: random (*H*_rand_), statistical bias (*H*_stat_) or deterministic rule (*H*_rule_). The mapping between left/right sides and statistics/rules was counterbalanced between subjects. Moreover, subjects answered several offline questions, notably about the moment when a regularity was detected. **c**, *Triangular arena*. The triangular arena provides a coordinate system where each location corresponds to a triplet of posterior probabilities of the three different hypotheses.

### Normative two-system model

We formalized our theoretical proposal as an ideal observer model of the task. This normative model combines the sequence of observations with prior knowledge about the task structure in order to constantly estimate, through Bayesian inference, the posterior probability of the three possible generative processes (i.e. random, random-to-statistical and random-to-rule). Under the random hypothesis (*H*_rand_), the model considers that observations are drawn randomly without bias. For the nonrandom hypotheses, the model considers that there is a single change-point that separates an initial random part from a second non-random part. On the one hand, for the random-to-statistical hypothesis (*H*_stat_), the model considers that the second part is characterised by a bias in the transition probabilities between successive observations. On the other hand, for the random-to-rule hypothesis (*H*_rule_), the model considers that the second part corresponds to the repetition of a particular pattern. The position of the change-point, the bias in transition probabilities, and the repeated pattern are unknowns that must be inferred from the sequence itself. Bayesian inference is used in all cases, providing a unified account of regularity detection independently of the type of regularity.

We simulated this model with the same sequences that were presented to subjects in order to derive quantitative predictions, and compare them to subjects’ behaviour. We also contrasted this *normative two-system model* to alternative models which we describe in more detail in the last section of the *Results*, and in *Methods*. In short, there are two types of alternative models: (1) *normative single-system models* use a single system, normatively, for both statistics and rules, instead of two hypothesis spaces; (2) *non-commensurable two-system models* use two systems, one per non-random hypotheses, but in a way that is not normative, rendering the two non-random hypotheses non commensurable.

### Offline, post-sequence reports

#### Generative process

At the end of each sequence, in order to assess subjects’ sensitivity in detecting either type of regularity, we asked them to retrospectively judge whether or not a regularity was present (detection) and, if so, to report its type (discrimination). Subjects were able to both detect the presence of a regularity and to discriminate its type (*Fig. 3a*; *Extended Data Fig. 2* for individual sequences): they performed above chance in identifying each of the three generative processes (on average 61.7%, 71.9%, and 90.4%; chance = 33.3%; *d*_Cohen_ ≥ 1.21, *t*_22_ ≥ 5.80, *p* ≤ 7.76 × 10^−6^). However, accuracy differed across sequence types (*w*^2^ = 0.022, *F*_2,22_ = 20.6, *p* = 4.86 × 10^−7^): subjects were better at detecting deterministic rules than statistical biases (difference in accuracy = 18.5%, CI = [11.4, 25.6], *d*_Cohen_ = 1.12, *t*_22_ =5.38, *p* = 2.12 × 10^−5^).

**Fig. 3.**
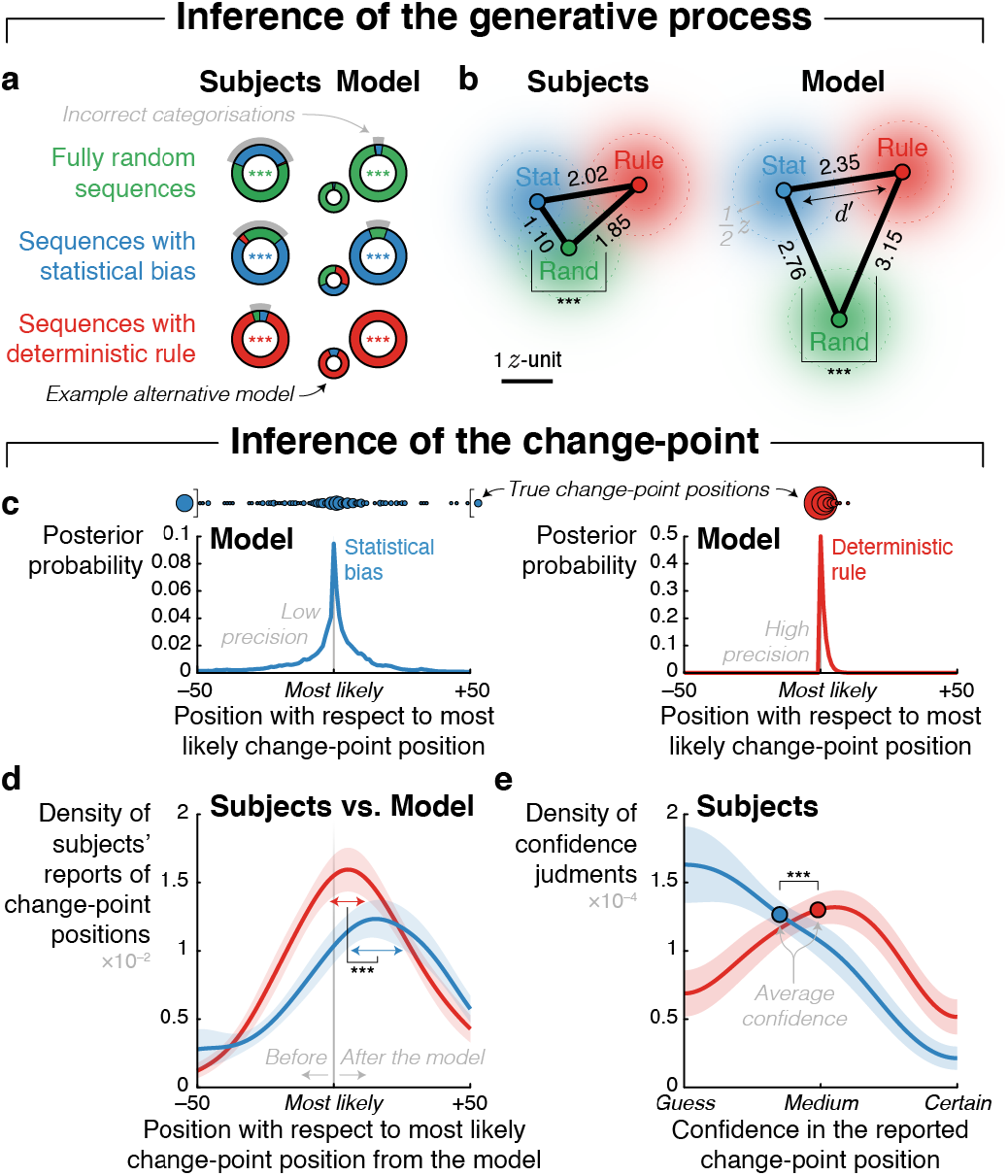
Offline (post-sequence) identification of the generative process and change-point position. **a**, *Post-sequence reports of sequence type*. Reported generative process is plotted as a function of the true generative process for the subjects and the *normative two-system mode* (simply “Model”). The categorisation profile depends on the underlying sequence processing as shown by the categorisation profile from an example alternative model, which detects both statistics and rules based on transition probability learning (of order 4) in all cases, and arbitrate between them based on predictability strength (see the *normative single-system model* in *Results’* last section for details). Models identify the most likely generative process based on the posterior probability of each hypothesis at the end of the sequence. **b**, *Signal detection theory analysis of sequence identification*. Multidimensional signal detection theory is applied on subjects reports in order to compare sensitivity to different regularities independently of possible response biases. Sensitivity measures (*d*’, in *z*-units) are represented as edges of a triangle whose vertices represent the three possible generative processes. Similar to the model, the subjects are worse at distinguishing statistical biases from a random process than they are for the deterministic rules. **c**, *Model beliefs about change-point position*. The posterior distribution over change-point positions (here, centred on the most likely position of the change-point as inferred by the model) reveals that the inference is more precise at detecting the onset of a deterministic rule than it is at detecting the onset of a statistical bias. **d**, *Subjects’ estimates of change-point position*. Subjects’ reports of change-points positions are aligned on the most likely change-point position as estimated by the model (as in **c**). Subjects are delayed relatively to the model, but, similarly to the model, their estimates are more precise for deterministic rules than statistical biases. Distributions reflect gaussian kernel densities with a bandwidth of 7 observations. **e**, *Subjects’ confidence in their estimates of change-point position*. Subjects reported higher confidence about change point for statistical biases than deterministic rules. Distributions reflect gaussian kernel densities with a bandwidth of 0.15 (where “guess” = 0 and “certain” = 1). The shaded area corresponds to the standard error of the mean. Stars denote significance: *** *p* < 0.005, ** *p* < 0.01, * *p* < 0.05.

To confirm that these results were not due to subjects’ preferring overall one category, and easily compare subjects and the *normative two-system model*, we relied on a (2D) signal detection theory analysis and quantified subjects’ sensitivities (*Fig. 3b*). As suggested by the previous analysis, the detection of a statistical bias was accompanied by a lower sensitivity than the detection of a deterministic rule (difference in *d’* = 0.76, CI = [0.55, 0.96], *d*_Cohen_ = 1.59, *t*_22_ = 7.62, *p* = 1.32 × 10^−7^). We applied the same signal detection theory analysis to the model, and the generative hypothesis that it estimates as most likely at the end of each sequence. As in subjects, the model’s sensitivity was lower for statistical biases compared to deterministic rules (difference in *d’* = 0.39, CI = [0.20, 0.59], *d*_Cohen_ = 0.89, *t*_22_ = 4.25, *p* = 3.31 × 10^−4^).

Interestingly, statistical biases that were missed by the subjects were not characterised by a late occurrence of the change-point (non-significant difference in change-point position = −2.84, CI = [−9.03, 3.35], *d*_Cohen_ = −0.20, *t*_22_ = −0.95, *p* = 0.35, BF_null_ = 3.05; *Extended Data Fig. 3a*) but instead by a lower posterior probability (i.e. closer to chance level) estimated by the model, relative to regularities that were detected (difference in *p*(*H*_stat_|seq) = 0.15, CI = [0.06, 0.25], *d*_Cohen_ = 0.68, *t*_22_ = 3.27, *p* = 0.004; *Extended Data Fig. 3b*). Subjects thus missed regularities more when the evidence was weaker. In particular, subjects more often missed alternation biases compared to both repetition biases (difference in detection rate = −23%, CI = [−36.4, −8.6], *d*_Cohen_ = −0.70, *t*_22_ = −3.36, *p* = 0.003) and frequency biases (difference in detection rate = −21%, CI = [−34.9, −6.4], *d*_Cohen_ = −0.63, *t*_22_ = −3.01, *p* = 0.006; *Extended Data Fig. 3c*). This reduced sensitivity to alternation is a cornerstone of transition probability learning ^23,54^.

Those categorization results are not a mere consequence of the task structure: the categorization profile of alternative models forced to do the same task is indeed different (*Fig. 3a* for an example alternative model and *Extended Data Fig. 2* for a more complete list). Although overall less accurate, subjects’ offline detection and identification of regularities were well accounted for by the *normative two-system model*, and better than alternative models (see *Results*’ last section).

#### Change-point position

Because of its hierarchical and normative nature, the model entertains a posterior distribution over possible change-point positions. Its maximum a posteriori estimate was highly correlated with the true change-point position across sequences, but less so in the case of statistical biases than for deterministic rules (difference in correlation *=* 0.53, CI = [0.38, 0.69], *d*_Cohen_ = 1.47, *t*_22_ = 7.05, *p* = 4.46 × 10^−7^; *Fig. 3c*), indicating that, even for a normative model, the detection of a change-point is more accurate for an emerging deterministic rule than for an emerging statistical bias. In order to test for the same effect in subjects, at the end of each sequence for which they reported the presence of regularity, we asked them to report the most likely position of the change-point, on a scale spanning the sequence length. We aligned subjects’ reports on the most likely change-point position as estimated by the model. Subjects’ reports were delayed compared to the model (statistical biases: delay = 7.40, CI = [2.51, 12.3], *d*_Cohen_ = 0.65, *t*_22_ = 3.14, *p* = 0.005; deterministic rules: delay = 7.47, CI = [3.39, 11.6], *d*_Cohen_ = 0.79, *t*_22_ = 3.79, *p* = 0.001; *Fig. 3d*). Importantly, as for the model, subjects’ reports were more precise in the case of deterministic rules compared to statistical biases (difference in variance = −1058.1, CI = [−1528.8, −587.3], *d*_Cohen_ = −0.97, *t*_22_ = −4.66, *p* = 1.20 × 10^−4^).

In order to probe whether subjects were aware that their estimates of the change-point position were less accurate for statistical biases than for deterministic rules, we asked them to report their confidence in their estimates. Reported confidence was indeed lower for statistics compared to rules (difference in confidence = 14.4%, CI = [11.0, 17.8], *d*_Cohen_ = 0.53, *t*_22_ = 8.76, *p* = 1.26 × 10^−8^). In the model too, the confidence about the change-point location, which can be formalized as the log-precision of the posterior distribution ^24,55,56^ over change-point positions, is also markedly lower for statistical regularities (difference in log-precision = 5.05, CI = [4.85, 5.25], *d*_Cohen_ = 11.1, *t*_22_ = 53.3, *p* = 9.46 × 10^−25^; *Fig. 3e*).

### Online report with finger tracking

We now turn to finger tracking recordings in order to explore whether subjects can faithfully track, in real time, the dynamics of their inference. Throughout the sequence, subjects continuously reported their current beliefs about the generative process of the observed sequence by moving their finger on a touch screen, within a triangular arena whose vertices correspond to each possible generative hypothesis: random, random-to-statistical, and random-to-rule (*Fig. 2c*). Importantly, each position in the triangle corresponds to a unique given set of posterior probability in the 3 possible hypotheses *H*_rand_, *H*_stat_, and *H*_rule_ (e.g. the bottom corresponds to (1, 0, 0), which is also the starting point, the centre to (⅓, ⅓, ⅓), etc.). Subjects’ finger trajectories are thus directly converted into a timeseries of posterior probabilities ascribed to each hypothesis, which can be compared to the timeseries of posterior probabilities estimated by the *normative two-system model (Fig. 4a,b*).

**Fig. 4.**
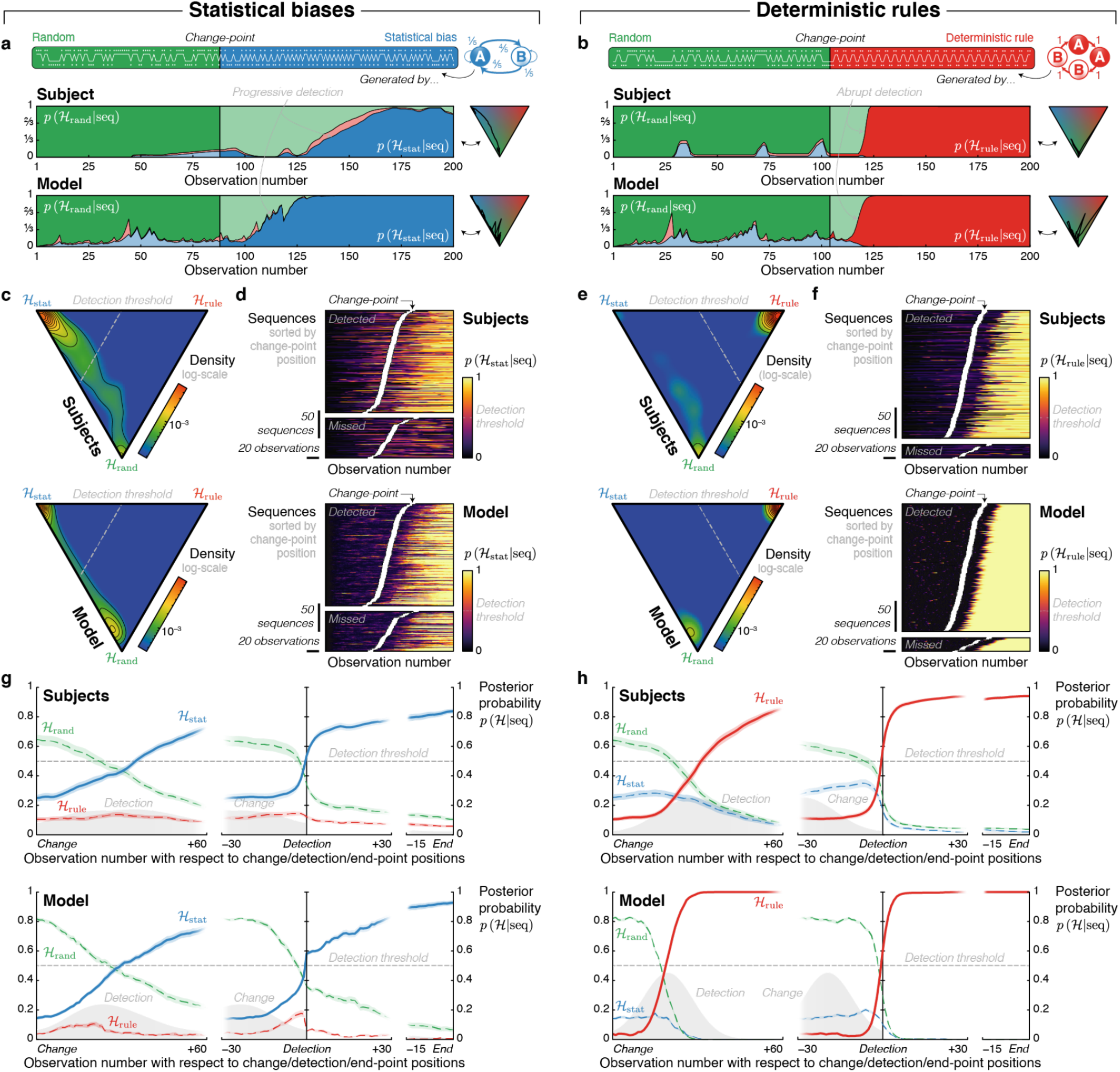
Different detection dynamics for statistical biases and deterministic rules. **a**/**b**, *Single-trial dynamics of hypothesis posterior probability* from an example subject and the model presented with the same random-to-statistical/rule sequence. **c**/**e**, *Triangular histograms* (smoothed and log-scaled) from the subjects and the model during parts of sequences with a statistical bias/deterministic rule. Different parts of the triangular arena are used in the different types of non-random parts of sequences. **d**/**f**, *Single-trial dynamics of the posterior probability in the statistical bias/deterministic rule hypothesis*. Sequences are grouped by detection (from subjects’ post-sequence reports) and sorted by change-point position. Note the progressive/abrupt detection of statistical bias/deterministic rule. **g**/**h**, *Averaged dynamics of posterior probability locked on change, detection and end points of parts of sequences with a statistical bias/deterministic rule*. The average increase in the probability of the statistical bias/deterministic rule hypothesis also shows the progressive/abrupt difference, in subjects and the model alike. This difference is even clearer when the curves are time-locked to the detection-point defined as the moment when *p*(*H*_correct_|seq) >½. Here, analyses were further restricted to non-random sequences that were correctly identified by subjects and for which detection-points were found for both the subjects and the model. Distributions reflect gaussian kernel densities of change-/detection-points with a bandwidth of 8 observations.

### Dynamics of regularity detection

Under the hypothesis that subjects approximate the *normative two-system model*, we expect the dynamics of regularity detection to be different in the case of statistical biases vs. deterministic rules. Below, we list those normative differences, illustrate them with the model and test them in subjects’ finger trajectories.

#### Overall distribution of beliefs

We created 2D triangular histograms of finger position across all sequences in order to quantify belief distributions. Subjects, like the model, used the bottom part of the triangle, corresponding to the random hypothesis, as well as the parts of the triangle close to the statistical bias hypothesis vertex during random-to-statistical sequences and, similarly, the parts of the triangle close to the deterministic rule hypothesis vertex in the random-to-rule sequences (*Fig. 4c,e*); thereby demonstrating a correct identification of the generative process in the course of the sequence.

#### Build-up of beliefs

In order to explore the differences in belief update for statistical biases and deterministic rules, we now turn to the analysis of time-resolved trajectories. In the model, the posterior probability of the correct hypothesis increased much more steeply in the case of rules than for statistics, resulting in step-like vs. gradual trajectories respectively. Subjects behave similarly, as can be seen on the trajectories locked on the true change-point position. However, locking on the true change-point underestimates the step-like nature of belief update in the case of deterministic rules due to variability in the onset of the step with respect to the change-point. We therefore also locked the trajectories onto the moment when the probability of the correct hypothesis becomes more likely than any other, which we refer to as the detection-point. This analysis revealed that in subjects, just as in the model, the increase in the posterior probability of the deterministic rule hypothesis is quite abrupt whereas the increase in the posterior probability of the statistical bias hypothesis is quite gradual (*Fig. 4g,h*). This difference is readily seen when inspecting individual trials (*Fig. 4d,f*). We quantified this difference by fitting sigmoid functions to trajectories in individual trials and averaged the fitted slope parameter across sequences: it was significantly higher for rules compared to statistics in the model (difference in slope = 0.39, CI = [0.33, 0.46], *d*_Cohen_ = 2.58, *t*_22_ = 12.4, *p* = 2.13 × 10^−11^) and in subjects (difference in slope = 0.27, CI = [0.18, 0.36], *d*_Cohen_ = 1.32, *t*_22_ = 6.35, *p* = 2.16 × 10^−6^; *Extended Data Fig. 4*).

### Regularity-specific modulations of detection dynamics

The *normative two-system model* predicts specific differences in detection dynamics within each type of regularity. More specifically, the strength of the statistical bias should modulate the amount of belief update, thus changing the slope of the gradual detection dynamics characteristic of statistical biases (*Fig. 5a*). By contrast, the length of the deterministic rule should change the moment of the detection-point, while maintaining the abrupt detection dynamics typical of deterministic rules (*Fig. 5b*).

**Fig. 5.**
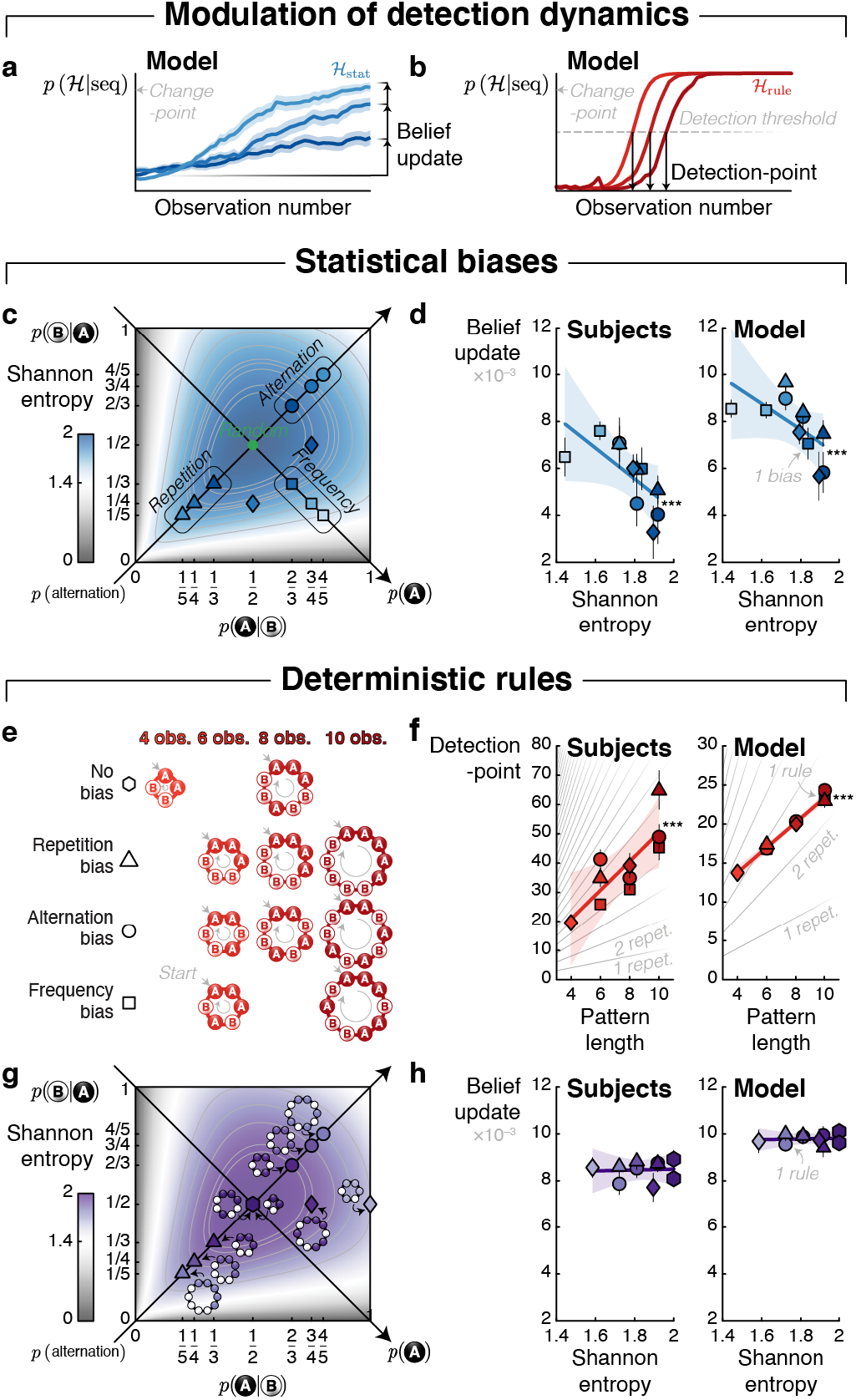
Regularity-specific, normative modulation of detection dynamics. **a**, *Examples belief updates from the model*. The amount of belief update, that is the difference in belief between the change-point and the end of the sequence, is expected to vary depending on the statistical bias considered. **b**, *Example detection-points from the model*. The amount of belief update is not expected to vary depending on the deterministic rule considered, only the detection-point, that is the number of observations between change- and detection-points (i.e. when the deterministic rule hypothesis is more likely than any other hypothesis). **c**, *Types of statistical biases*. Statistical biases are defined by first-order transition probabilities, which have different subtypes of bias (item frequency, repetition-alternation frequency) and strength (measured by Shannon entropy). **d**, *Modulation of belief update for statistical biases*. Change-points, if detected, should lead to belief update. In random-to-statistical sequences, the amount of belief update decreased with Shannon entropy of generative first-order transition probabilities. **e**, *Types of deterministic rules*. Deterministic rules are defined by the pattern that is repeated, which has a specific length, and which induces a specific statistical bias. **f**, *Modulation of detection-point for deterministic rules*.Change-points, if detected, should lead to crossing of the detection threshold. In random-to-rule sequences, the detection-point increased with pattern length. **g**, *Apparent statistical biases of deterministic rules*.Repeating patterns induce, on purpose, different types of statistical biases, of different strength (measured as Shannon entropy). Most of them are matched with biases induced by pure statistical biases. **h**, *No modulation of belief update for deterministic rules*. Contrary to pure statistical biases, belief update of deterministic rules does not decrease with Shannon entropy of apparent first-order transition probabilities. Stars denote significance: *** *p* < 0.005, ** *p* < 0.01, * *p* < 0.05.

#### Statistical biases

We sorted statistical biases according to the type and amount of bias they induce: one item could be more frequent than the other, or repetitions could be more (or less) frequent than alternations, or both of these two biases could be present. We quantified the strength of this bias by the Shannon entropy of the corresponding generative transition probabilities, which ranged from 1.44 to 1.92 bit here (*Fig. 5c*). All statistical biases were detected gradually, but the amount of belief update varied across regularities (*Fig. 5d*): with a significant effect of entropy for the model (correlation = −0.26, CI = [−0.42, 0.10], *d*_Cohen_ = −0.71, *t*_22_ = −3.40, *p* = 0.003) and the subjects (correlation = −0.31, CI = [−0.47, 0.14], *d*_Cohen_ = −0.81, *t*_22_ = −3.89, *p* = 7.89 × 10^−4^).

#### Deterministic rules

We distinguished among deterministic rules according to their lengths (4, 6, 8 or 10 observations; *Fig. 5e*). Furthermore, some of the patterns purposely comprised a bias in the apparent transition probabilities between items (*Fig. 5g*). For instance, the repetition of the AAAABBBB pattern results in many more repetitions (75%) than expected from a random process (50%). By contrast, the repetition of AABB produces as many As and Bs (50%) and as many repetitions as alternations (50%), which is equal to what is expected on average from a random process. The results showed that the detection-point (*Fig. 5f*) increased with the length of the pattern for both the model (correlation = 0.87, CI = [0.79, 0.95], *d*_Cohen_ = 4.48, *t*_22_ = 21.5, *p* = 2.99 × 10^−16^) and the subjects (correlation = 0.51, CI = [0.43, 0.59], *d*_Cohen_ = 2.76, *t*_22_ = 13.3, *p* = 5.80 × 10^−12^). Importantly, even though the detection-point varied according to the length of the patterns, the detection dynamics always remained similarly abrupt (*Fig. 5h*); in particular, belief update was not modulated by the strength of the statistical bias induced by the pattern (quantified by Shannon entropy), neither in the model (non-significant correlation = 0.075, CI = [−0.083, 0.23], *d*_Cohen_ = 0.21, *t*_22_ = 0.99, *p* = 0.36, BFnull = 2.96) nor in subjects (non-significant correlation = 0.080, CI = [–0.069, 0.23], *d*_Cohen_ = 0.23, *t*_22_ = 1.11, *p* = 0.28, BFnull = 2.64).

### False alarms in random sequences

Throughout random sequences (i.e. the initial part of random-to-non-random sequences and the entire length of random sequences), the *normative two-system model* predicts that observers keep track of all hypotheses. Thus, they should entertain high probabilities in the random hypothesis, as indeed observed; but because random sequences contain transient periods of apparent regularity, beliefs in non-random hypotheses should also occasionally increase (*Fig. 6a*). Results indicate that such false alarms do occur in subjects. Furthermore, in both the model and the subjects, they much more often concerned a transient preference for the statistical bias hypothesis than for the deterministic rule hypothesis (*Fig. 6b*). This is because local trends, even weak, constantly impact the statistical hypothesis by slightly shifting the estimated probabilities away from chance. By contrast, transient trends impact the deterministic hypothesis only when they license strong (erroneous) predictions, which rarely occur by chance. Below, we explore the dynamics and rationality of subjects’ false alarms during random sequences.

**Fig. 6.**
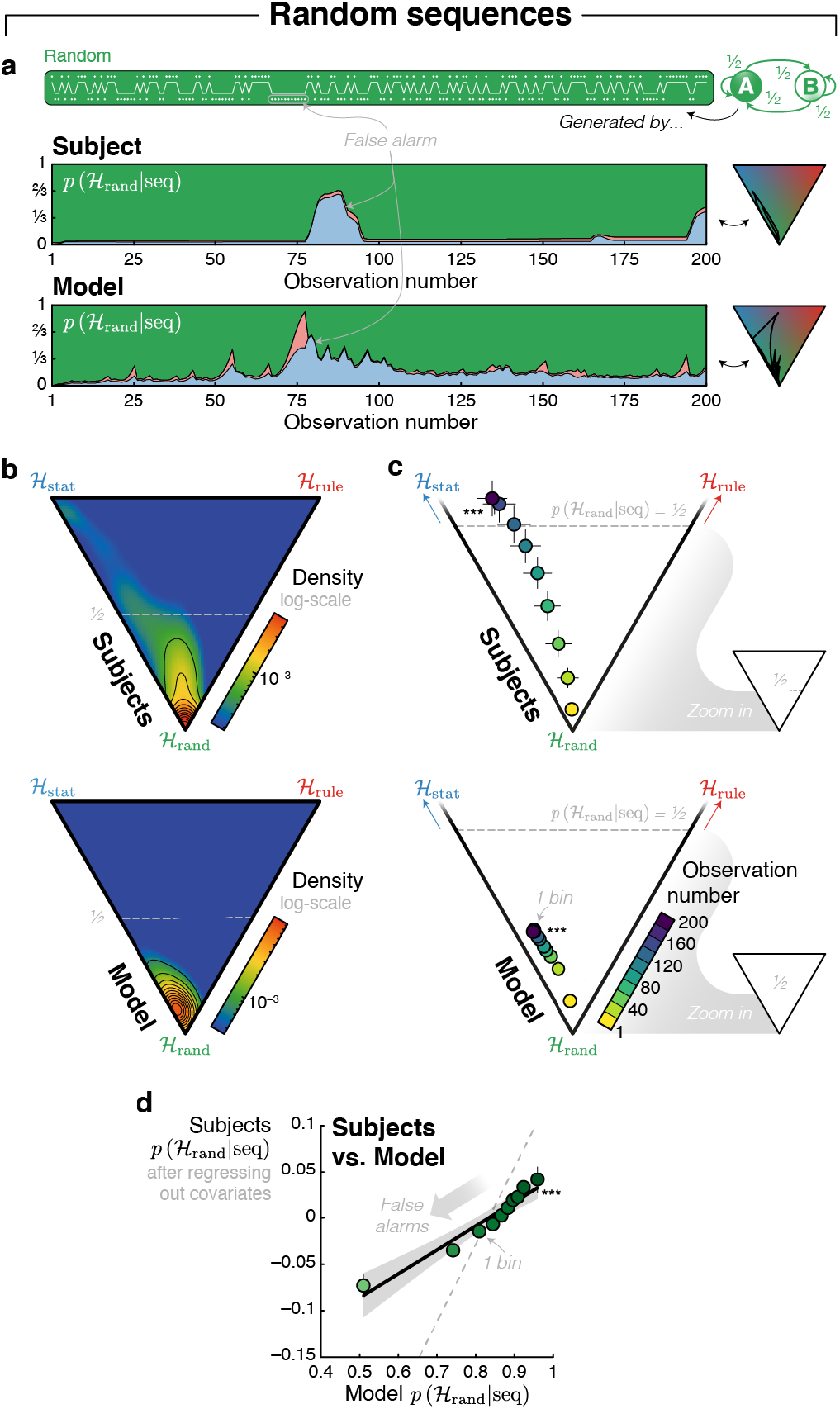
False alarms reflect transient periods of regularity. **a**, *Example random sequence and resulting inferences*. Transient periods of regularity can appear, by chance, in random sequences (i.e. the initial part of random-to-non-random sequences, or the entire length of the random sequences). They can lead observers to make false alarms, that is, decrease the posterior probability of the random hypothesis in favour of a non-random hypothesis. **b**, *Triangular histograms* (smoothed and log-scaled) for the subjects (top panel) and the model (bottom panel) during random sequences. **c**, *Effect of observation position within the sequence on false alarms*. Reported probabilities from the subjects and the model are averaged in groups of 20 consecutive observations in the sequence. The false alarm rate increased as more observations were received within a sequence, in subjects (top panel) and the model (bottom panel). **d**, *Correlation between subjects’ and model’s posterior probability in the random hypothesis*. A linear relationship between subjects and the model’s probability of the random hypothesis (in 10 bins defined using deciles) remains even after regressing-out the confounding (normative) effect of elapsed time in the sequence from subjects’ data (dashed line: without regressing-out). Stars denote significance: *** *p* < 0.005, ** *p* < 0.01, * *p* < 0.05.

We first tested whether subjects’ false alarms increase with the number of observations received (*Fig. 6c*). This is a prediction of the model: the posterior probability of the random hypothesis steadily decreases with the number of observations in the sequence (correlation = −0.32, CI = [−0.34, −0.29], *d*_Cohen_ = −5.53, *t*_22_ = −26.5, *p* = 3.45 × 10^−18^). This is a cornerstone of hierarchical inference in our task: the model assumes that a change-point occurs in about two thirds of sequences, which translates into a slowly increasing probability that a change-point has occurred as the sequence unfolds in time. Subjects show the same effect (correlation = −0.60, CI = [−0.67, −0.54], *d*_Cohen_ = −4.18, *t*_22_ = −20.1, *p* = 1.24 × 10^−15^).

We then tested whether false alarms occurred precisely when the model predicted them, that is when the random sequences exhibited, by chance, a transient regularity. To do so, posterior probabilities in the random hypothesis reported by subjects were regressed against the model’s posterior probabilities in the random hypothesis (*Fig. 6d*). We found a significant positive effect (coefficient = 0.58, CI = [0.50, 0.65], *d*_Cohen_ = 3.34, *t*_22_ = 16.0, *p* = 1.27 × 10^−16^) that also survived when the number of observations received was included as covariate (coefficient = 0.23, CI = [0.16, 0.30], *d*_Cohen_ = 1.41, *t*_22_ = 6.77, *p* = 8.41 × 10^−7^). This result confirms that subjects’ false alarms reflect a normative inference due to transient deviations from randomness, over and above the simple effect of elapsed time since the beginning of the sequence.

### Normative graded weighting of non-random hypotheses

A crucial assumption of the *normative two-system model* is that subjects perform a Bayesian inference over two distinct hypothesis spaces in order to detect and identify statistical biases and deterministic rules. The normative nature of this inference enables to compute the posterior probability of the different non-random hypotheses in the same probabilistic currency and, therefore, to compare them. To study this prediction, we now explore situations of conflict, when both non-random hypotheses compete without a clear winner. Those situations appear frequently just before the detection of a deterministic rule as well as during random sequences. We used the belief difference between non-random hypotheses (i.e. the difference between their posterior probability normalized by their sum) as an index of conflict between statistics and rules in those periods.

*Before the detection of deterministic rules*, the model often shows a period of indecision during which nonrandom hypotheses have intermediate posterior probabilities. Those intermediate levels temporarily favour the statistical bias hypothesis before a change-of-mind ^57^ leads to an accurate identification of the deterministic rule, and does so all the more that the repeated pattern has a strong bias in its apparent transition probabilities (*Fig. 7a*). To assess this effect, we correlated the strength of this bias (quantified by Shannon entropy) with the belief difference averaged during the period going from the true change-point to the detection-point in the model (correlation = 0.49, CI = [0.37, 0.61], *d*_Cohen_ =1.70, *t*_22_ = 8.17, *p* = 4.14 × 10^−8^) and in the subjects (correlation = 0.19, CI = [0.0013, 0.37], *d*_Cohen_ = 0.44, *t*_22_ = 2.09, *p* = 0.049). Importantly, the belief difference assessed in the model and in subjects’ data during this indecision period correlated with each other across patterns (correlation = 0.22, CI = [0.078, 0.37], *d*_Cohen_ = 0.67, *t*_22_ = 3.20, *p* = 0.004; *Fig. 7b*).

**Fig. 7.**
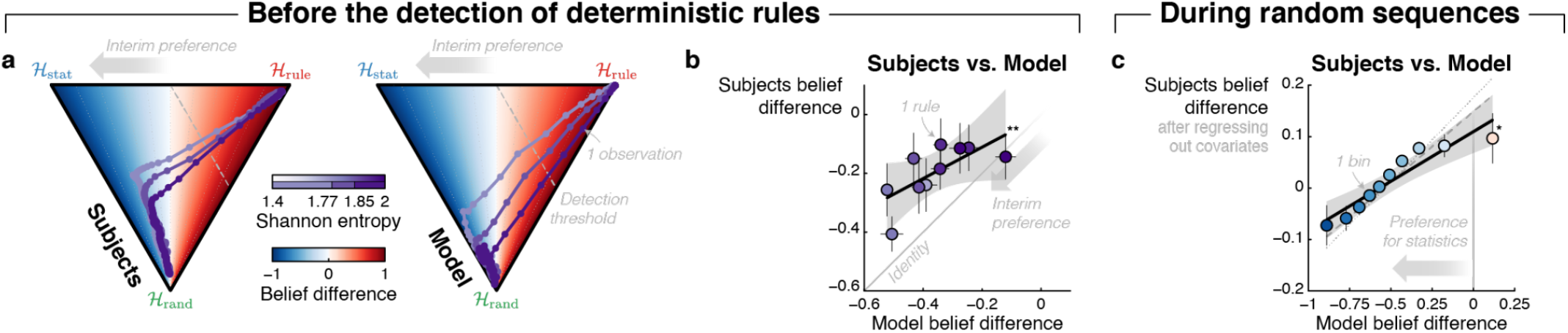
Normative graded weighting of non-random hypotheses. **a**, *Interim preference for the statistical bias hypothesis before the detection of deterministic rules depends upon the entropy of the apparent transition probabilities characterising the pattern*. Trajectories during random-to-rule sequences were centred on the detection-point and binned according to the entropy of the repeated pattern. Each dot corresponds to one observation in the sequence. The belief difference is the difference between the posterior probability of the two non-random hypotheses normalized by their sum (it evolves between –1 and 1 independently of the probability of the random hypothesis), which is shown here as a heatmap. **b**, *Belief difference from subjects as a function of the belief difference from the model across patterns*. The belief difference between non-random hypotheses was averaged from the change-point to the detection-point, in both subjects and the model. Each dot corresponds to one rule, whose entropy is color-coded. **c**, *Belief difference from subjects as a function of the belief difference from the model in random sequences*. In random sequences (i.e. the initial part of random-to-non-random sequences, or the entire length of the random sequences), the belief difference from subjects was regressed against the model belief difference. The linear relationship between subjects and the model belief difference is shown (in 10 bins defined using deciles) after regressing-out effect of confounding variables such as the posterior probability of the random hypothesis (dotted line: without regressing-out), and together with its interaction with the posterior difference in the regression (dashed line: without regressing-out). Stars denote significance: *** *p* < 0.005, ** *p* < 0.01, * *p* < 0.05.

*During random sequences*, while the model generally assigns a low probability to both statistics and rule hypotheses, their relative posterior probability fluctuates depending on the exact observations received. We reasoned that if subjects used a normative arbitration between non-random hypotheses, the relative credence they assign to both hypotheses should show fluctuations similar to the model. We therefore regressed subjects’ difference in beliefs between non-random hypotheses against the model’s difference in beliefs (*Fig. 7c*). We found a significant positive effect (coefficient = 0.28, CI = [0.21, 0.35], *d*_Cohen_ = 1.71, *t*_22_ = 8.18, *p* = 4.08 × 10^−8^) that also survived when confounding variables were included in the regression model, including the posterior probability of the random hypothesis, alone (coefficient for belief difference = 0.24, CI = [0.18, 0.31], *d*_Cohen_ = 1.55, *t*_22_ = 7.41, *p* = 2.04 × 10^−7^) or together with its interaction with the belief difference (coefficient for belief difference = 0.17, CI = [0.02, 0.33], *d*_Cohen_ = 0.49, *t*_22_ = 2.33, *p* = 0.029).

### Alternative models

The *normative two-system model* accounts for many aspects of the human detection and identification of temporal regularities observed in the current experiment. To test the necessity of the model’s assumptions, we explored alternative models lacking one or the other of its assumptions. We tested, first, models with a single common system for deterministic rules and statistical biases (instead of two hypothesis spaces) and second, models with distinct but non-commensurable systems (instead of the shared, normative probabilistic currency). We evaluated those models on three features of subjects’ behaviour: the categorisation of sequences, the detection dynamics of deterministic rules, and the comparison of non-random hypotheses during random sequences (*Extended Data Fig. 5*).

#### Normative single-system model

This model posits that subjects use a continuum rather than two distinct hypothesis spaces for statistics and rules (hence *single-system* instead of *two-system):* because deterministic rules are a limit case of statistical biases (with *p* = 0 or 1), subjects could employ statistical learning in all cases. This is because the repetition of a pattern necessarily induces a bias in the apparent higher-order transition probabilities (e.g. repetitions of AABB result in a bias in order-2 transition probabilities: *p*(B|AA) = *p*(A|BA) = *p*(B|AB) = *p*(A|BB) = 1).

We tested different versions of the *single-system model* in a factorial fashion (*Fig. 8a*). We started by tying *H’*_rule_ to the monitoring of higher-order transition probabilities (we tested orders 2 to 9), while keeping *H*_stat_ tied to the monitoring of first-order transitions as before. In terms of sequence categorisation, with orders smaller than 3, the *single-system model* missed some repeated patterns, and for higher orders, it increasingly miscategorized fully random sequences as random-to-rule ones (*Fig. 8b* and *Extended Data Fig. 2*) such that, no matter the order used, it always provided a worse account of subjects’ categorisation than the *two-system model* (relative MSE ≥ 0.018, *d*_Cohen_ ≥ 0.67, *t*_22_ ≥ 3.22, *p* ≤ 0.004; *Fig. 8c*). In terms of detection dynamics, we previously investigated their progressive vs. abrupt nature (*Fig. 4*), a feature that only models with order larger than 4 also exhibited (*Extended Data Fig. 6*). However here, we focused on a simpler feature: while the detection dynamics of rules in the *normative two-system model* and subjects is smooth, in the *single-system model* it appears unsteady, repeatedly switching between acceleration and deceleration (which we quantified as the average absolute acceleration, i.e. second-order derivative: see *Methods* and *Extended Data Fig. 5b*). Specifically, in this version of the *single-system model*, those unsteady dynamics (see the staircase-like dynamics in *Fig. 8d*) led to a poor account of subjects’ behaviour independently of the transition order considered (relative MSE ≥ 2.35 × 10^−5^, *d*_Cohen_ ≥ 0.79, *t*_22_ ≥ 3.77, *p* ≤ 0.001; *Fig. 8e*).

**Fig. 8.**
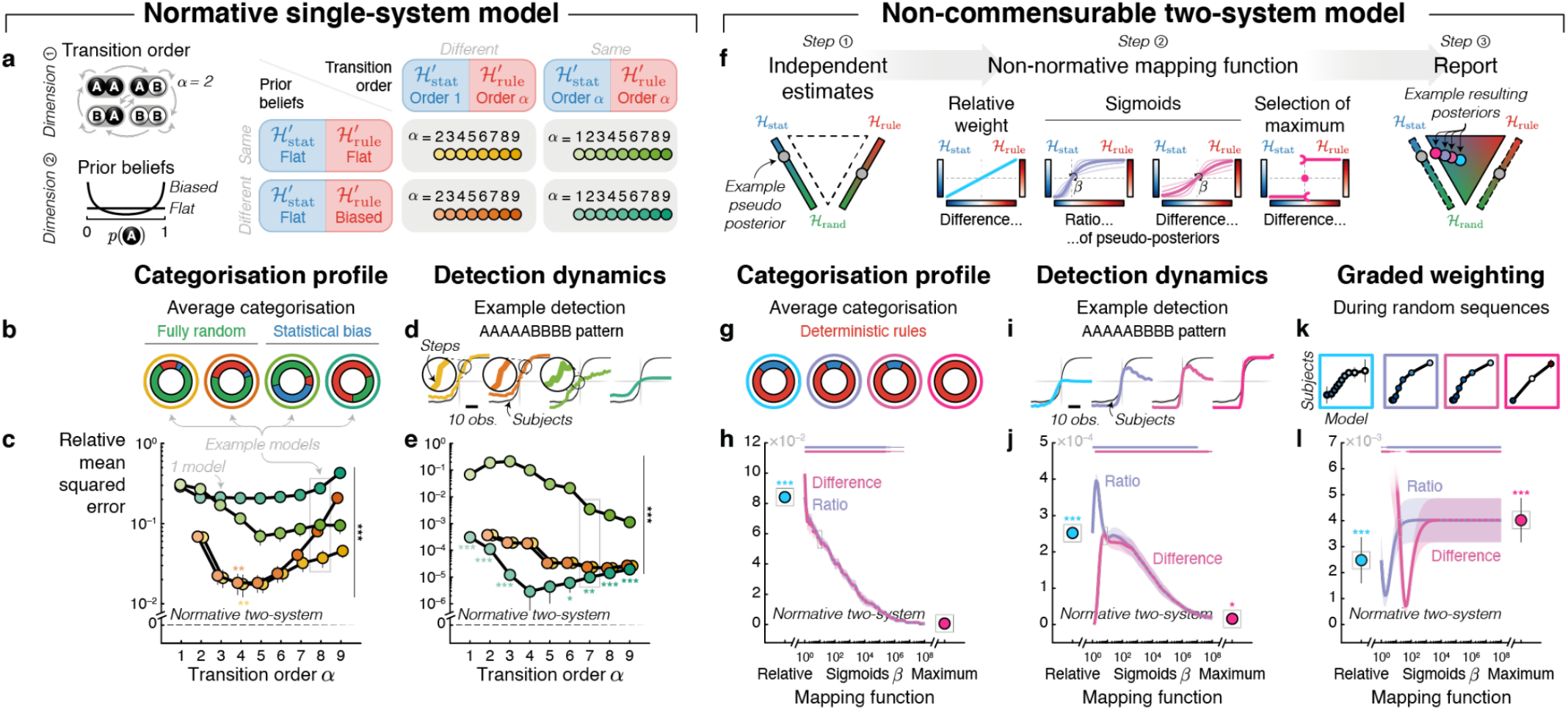
Evidence against alternative models. **a**, *Normative single-system model and its variants*. This alternative model uses statistical learning in all cases, to detect both statistical biases and deterministic rules. We varied in a factorial manner the order of transition probabilities that are estimated (*α*, from 1 to 9) and the prior beliefs about predictability (*d*, flat or biased), giving rise to 4 families of models. When *H*_stat_ and *H*_rule_ correspond exactly to the same hypothesis (same transition order, flat priors; upper-right cell), we used the strength of predictions to arbitrate between them. Otherwise, the arbitration was based on the posterior probabilities of hypotheses, as in the *normative two-system model*. **b**/**g**, *Example categorisation profiles from alternative models;* to be compared with *Fig. 3a*. **c**/**h**, *Error of alternative models with respect to subjects’ categorisation profiles*. The mean squared error (MSE) is measured using the error matrix presented in *Extended Data Fig. 5a*. **d**/**i**, *Example detection dynamics of the AAAAABBBBB deterministic rule by alternative models*. The detection dynamics are locked on detection-points; to be compared with *Fig. 4h*. The black curve corresponds to subjects. **e**/**j**, *Error of alternative models with respect to subjects’ detection dynamics of deterministic rules*. The mean squared error (MSE) is measured on the absolute acceleration (average absolute second-order derivative) of the posterior probability of the deterministic rule after the change point, as presented in *Extended Data Fig. 5b*. **f**, *Non-commensurable two-system model*. This alternative model uses the same three hypotheses as the *normative two-system model* but it estimates the (pseudo) posterior probability of each non-random hypothesis independently from one another. It then combines non-normatively those (pseudo) posterior probabilities to map them onto the triangle. We explored several combination functions: linear, maximum, and sigmoid functions (of which we varied the slope, *β*) on the difference or on the ratio of (pseudo) posteriors (see the example posteriors in step #3). k, *Example fit of subjects’ graded weighing of non-random hypotheses by alternative models*. Subjects’ belief difference in random sequences is plotted against the model’s; to be compared with *Fig. 7c*. **l**, *Error of alternative models with respect to subjects’ graded weighing of non-random hypotheses*. The mean squared error (MSE) is the residual error of the linear regression relating subjects’ belief difference in random sequences to the models’, restrained to ≥ 0 regression coefficients to prevent anti-correlation between models and subject, as presented in *Extended Data Fig. 5c*. In **c**, **e**, **h**, **j**, and *i*, MSE is plotted relatively to the *normative two-system model*; positive values indicate that the alternative models perform worse. Stars and line thickness above **h**/**j**/**l** denote significance: *** *p* < 0.005, ** *p* < 0.01, * *p* < 0.05. The examples can be changed with the online code, here we used *α* = 8 in **b**, *α* = 7 in **d**, *β* = 100 in **g**, *β* = 6 in **i**, *β* = 20 in **k**.

Next, we considered the possibility that *H’*_rule_ could not only be tied to the monitoring of higher-order transitions, but also to a prior belief biasing the inference towards predictability (i.e. inferred probability of the next item closer to 0 and 1). In this second version, *H*_stat_ remains tied to first-order transition probabilities with a flat prior, as before. This version performed worse than the previous one as soon as the slightest bias was introduced in *H’*_rule_ (*Fig. 8b-e* shows the results for *d* = 0.001; *d* = 0 is the flat prior; see *Methods*) and even worse for stronger priors (as can be checked with the online code).

In a third version, *H’*_rule_ and *H’*_stat_ were tied to the monitoring of the *same* transition order (we tested orders 2 to 9) but with a *different* prior, biasing the inference toward predictability in *H’*_rule_ (*d* > 0), but not in *H’*_stat_ (*d* = 0, flat prior). Because of such bias, many random-to-statistical sequences were miscategorised as random-to-rule (*Fig. 8b* and *Extended Data Fig. 2*), leading to an overall poorer account of subjects’ categorization (relative MSE ≥ 0.21, *d*_Cohen_ ≥ 2.63, *t*_22_ ≥ 12.61, *p* ≤ 1.52 × 10^−11^; *Fig. 8c*). The bias did not change much the detection dynamics of rules: subjects’ data remained accounted significantly better for by the *two-system model* for orders 1 to 3 and 7 to 9 (relative MSE ≥ 9.77 × 10^−6^, *d*_Cohen_ ≥ 0.60, *t*_22_ ≥ 2.87, *p* ≤ 0.009), and non-significantly for orders 4 to 6 (relative MSE ≥ 2.86 × 10^−6^, *d*_Cohen_ ≥ 0.26, *t*_22_ ≥ 1.24, *p* ≤ 0.23; *Fig. 8e*). The online code enables one to check that those results obtained with *d* = 0.001 become worse with larger values.

The last cell of the factorial model space is that *H’*_stat_ and *H’*_rule_ are tied to the monitoring of the *same* transition order (we tested 1 to 9) with the *same* flat prior. If *H’*_stat_ and *H’*_rule_ are the same, then normative categorization is impossible; we nevertheless explored the following hack: one weighs non-random hypotheses based on predictability, tying rules and statistics to higher and lower predictability, respectively. No matter the transition order used, this version poorly accounted for the subjects’ categorization (relative MSE ≥ 0.069, *d*_Cohen_ ≥ 0.89, *t*_22_ ≥ 4.28, *p* ≤ 3.02 × 10^−4^; *Fig. 8c*) and the smooth detection dynamics found in subjects for rules (relative MSE ≥ 0.0011, *d*_Cohen_ ≥ 1.18, *t*_22_ ≥ 5.67, *p* ≤ 1.05 × 10^−5^; *Fig. 8e*) compared to the *two-system model*. Those results were obtained when mapping linearly non-random hypotheses onto predictability, non-linear functions did not provide a better fit than the *two-system model (Extended Data Fig 7*).

#### Non-commensurable two-system model

This model considers the three hypotheses together (i.e. *H*_rule_, *H*_stat_, and *H*_rand_) and assigns to each of them a posterior probability that takes into account the other two hypotheses (through normalization in Bayes’ rule). A second alternative to this *normative model* is that subjects use distinct hypothesis spaces (i.e. *H*_rule_, *H*_stat_, and *H*_rand_ as before) but estimate the posterior probabilities of each non-random hypothesis only relatively to the random hypothesis, while ignoring the third hypothesis (i.e.*H*_rule_ vs. *H*_rand_ and *H*_stat_ vs. *H*_rand_). In this case, owing to a lack of normalization across the three hypotheses, their probabilities do not sum to 1 and cannot be directly compared (hence *non-commensurable* instead of *normative*), we thus term those probabilities ‘pseudo posteriors’. The *non-commensurable model* must resort to a non-normative way of comparing hypotheses in order to report a belief in the triangular arena; we explored several ways.

A first version of this *non-commensurable model* looks for the maximum: (1) it selects among the two putative regularities based on the maximum pseudo posteriors, and (2) it selects between the selected regularity and the random hypothesis according to the pseudo posterior. Regarding categorisation profiles, this version is indistinguishable from the *normative two-system model (Fig. 8g,h*). However, it produces different detection dynamics of deterministic regularities (*Fig. 8i*), which are so abrupt that they become discontinuous, resulting in a worse account of subjects’ data compared to the *normative model* (relative MSE = 1.59 × 10^−5^, *d*_Cohen_ = 0.47, *t*_22_ = 2.25, *p* = 0.035; *Fig. 8j*). In addition, because this model compares between the non-random hypotheses in a categorical rather than continuous manner, it cannot reproduce the graded weighting of non-random hypotheses found in subjects (*Fig. 7c*) and in the *normative model* during random sequences (relative MSE = 0.0025, *d*_Cohen_ = 0.99, *t*_22_ = 4.75, *p* = 9.76 × 10^−5^; *Fig. 8k,l*).

A second version of this *non-commensurable model* weighs the two non-random hypotheses according to the size of the difference in their pseudo posterior probabilities, rather than selecting one in an all-or-none manner. The problem with this version is that it often remains undecided, converging to a conclusion only if one of the two pseudo posterior probabilities vanishes (i.e. becomes 0). As a result, it cannot detect deterministic rules that have an apparent bias in transition probabilities (e.g. AAAAABBBBB; left panel of *Fig. 8i*), leading to a worse account of subjects’ categorisation profiles (relative MSE = 0.084, *d*_Cohen_ = 5.23, *t*_22_ = 25.1, *p* = 1.13 × 10^−17^; *Fig. 8g,h*) compared to the *normative model*. It is also worse regarding the detection dynamics of rules (relative MSE = 2.52 × 10^−4^, *d*_Cohen_ = 3.34, *t*_22_ = 16.0, *p* = 1.29 × 10^−13^; *Fig. 8i,j*). Because it often remains too undecided, this model also provided a worse account of the subjects’ graded weighting of non-random hypotheses during random sequences (relative MSE = 0.0025, *d*_Cohen_ = 0.59, *t*_22_ = 2.82, *p* = 0.010; *Fig. 8k,l*).

As an attempt to overcome the indecision problem, we tested a third version of the *non-commensurable two-system model* that can exaggerate the difference in pseudo posterior probabilities of the non-random hypotheses. We used a sigmoid function to formalize such exaggeration because it allows a significant impact of even subtle differences in pseudo posterior probabilities. Varying the slope of this sigmoid actually moves the model along a continuum whose ends correspond to the ‘maximum selection’ and the ‘linear difference’ versions tested above. However, this version suffered from the same problems as these previous versions (*Fig. 8g,i,k*) and no matter its parameterization, it always provided a worse account of subjects’ behaviour compared to the *normative model (Fig. 8h,j,l*). Note finally that resorting to a sigmoid that uses the log-ratio of (instead of the difference in) the pseudo posterior probabilities of the non-random hypotheses leads to the same conclusions.

#### Summary

Most alternative models performed worse than the *normative two-system model* on every feature we tested. At best, some alternative models appeared statistically indistinguishable from the *normative two-system model* on one feature, but they always performed worse on other features (*Extended Data Fig. 7* for all simulations). The *normative two-system model* therefore provided a better account of human behaviour than those many different alternative models. This finding held even though most of those alternative models had free parameters (e.g. transition order, strength of prior beliefs, sigmoid parameter, etc.) that gave more flexibility than the parameter-free *normative two-system model*.

## Discussion

We proposed here that a simple taxonomy may help to organise the theoretically infinite number of temporal regularities that exist in the environment. This taxonomy distinguishes deterministic rules, for which certainty can be attained, from statistical biases, for which a degree of uncertainty remains. Here, we asked human subjects to detect and identify regularities of either category that could suddenly appear within a random sequence of binary observations. We measured the dynamics of their inference by continuously tracking subjects’ finger movement ^57,58^ and compared them to normative models ^59,60^. We found that subjects assigned probabilistic credence to each of three distinct hypotheses, and that they treated statistics and rules as distinct hypothesis spaces rather than as a continuum, following our taxonomy.

Many previous studies focused on how humans use past observations to make predictions about the future. Those predictions leverage, often in different studies, either the estimation of a particular statistics (e.g. item frequency and transition probabilities) or the identification of a rule (e.g. a repeating pattern). We would like to stress that, in the context of predictions, the *characterization* of a specific generative process (estimation of a statistical bias, or the identification of a specific deterministic rule) is a different computational problem than the *identification* of the most relevant process (rule or statistics?) for predictions. The latter has received much less attention than the former, and it is the problem we tackled here. In the past, the detection of regularity (random vs. non-random) has indeed received a lot of attention. Notably, several studies on the subjective perception of randomness asked human subjects to categorize sequences as random versus non-random, and described human randomness perception in terms of statistical inference ^54,61–63^, which is in line with the theoretical account presented here. In another line of research, rule learning paradigms also prompted subjects to categorize random vs. non-random sequences ^10^. In particular, a recent study ^43^ found that this categorisation relies on the computation of a posterior distribution over possible patterns (of different lengths), a finding we also replicated here, while also extending it to other generative processes.

An innovation of our work is to study the advent of regularity. Real-world environments are usually volatile, causing frequent changes in the process generating the observations. As a consequence, in addition to categorising among regularities, observers trying to predict future observations must also detect *when* regularity appears. We found that both statistics and rules could be promptly detected, but with different detection dynamics. Deterministic rules were detected quite abruptly, reflecting ‘aha moments’^64^; which is a computational consequence of the all-or-none predictions they afford. The onset of the detection varied as a function of pattern length. Both findings replicate previous studies in which repeating patterns appeared suddenly in otherwise random sequences ^10,11^. However, it remains unclear whether subjects detected deterministic rules by monitoring the exact pattern being repeated or rather by detecting a subpart of it forming a noticeable temporal compound (e.g. the alternating series embedded in A<u>ABABABABB</u>), a well-known ability of the auditory system ^65,66^. By contrast to deterministic rules, statistical biases were detected more progressively, reflecting evidence accumulation ^28,57,67^; which is a computational consequence of their inherent uncertainty. The slope of the detection varied as a function of the strength of the bias (quantified in terms of Shannon entropy). Again, both findings replicate previous studies ^12,14^.

Detecting the onset of a regularity, be it deterministic or statistical, requires observers to explicitly monitor change-points separating random from non-random observations. Because change-point positions are a priori unknown, they must be inferred from the sequence, and the uncertainty associated with this inference of position should be factored into the evaluation of the non-random hypotheses, which is a form of hierarchical inference. Here, given that a change-point occurred in two thirds of sequences, the a priori probability of having encountered a change-point increased as the sequence unfolded. Accordingly, in random sequences, subjects’ false alarms increased with sequence duration, as also observed in a recent study ^12^. A model lacking such an explicit representation of change-points would, by contrast, have a stable false alarm rate throughout the sequence ^68^.

Another consequence of assuming the existence of change-points is the progressive down-weighting of past observations (*Supplementary Note 2*). This is because pre-change-point observations are discarded from the inference whenever change-points are suspected. Marginalising over several different possible change-points positions (with different posterior probabilities) thus progressively discounts previous observations. Such progressive discounting of previous observations is also observed in models with a memory decay ^22,23,54,69,70^ although the origin of the phenomenon is computationally different. Importantly, subjects’ behaviour is compatible with such a progressive discounting of past observations, which the *normative two-system model* thus automatically reproduces.

Subjects were also able to explicitly report change-point positions after sequence presentation, and rate their confidence in that estimation, both in a manner conforming to a hierarchical inference. This replicates the findings of recent studies on statistical learning and the explicit representation of changepoints ^20,22,24,25,71^; but extended here to the case of deterministic rules.

Our results indicate that the human detection of regularity and categorisation between regularities rely on two systems: one for detecting statistical biases and one for detecting deterministic rules, which we formalized as the *normative two-system model*. We wish to dissipate a potential misunderstanding: subjects’ reliance on a two-system model was not imposed by our three-alternative categorization task. As we explored in the *normative single-system model*, they could have solved this task using a single generative model (i.e. statistical learning), possibly with different parameters, for both statistics and rules. According to the *single-system model*, statistics and rules are regularities that vary in degree, while according to the *two-system model*,they belong to different hypothesis spaces. Measures as coarse as the profile of sequence categorisation suffice to tease apart the *normative single-* vs. *two-system models*. Moreover, during online inference, these two models also make different predictions regarding the graded weighing of non-random hypotheses and the detection dynamics of deterministic rules. Comparison between the two models revealed that subjects treated statistics and rules as pertaining to fundamentally different hypothesis spaces.

A question for future research is whether the distinct hypothesis spaces for statistics vs. rules are supported by distinct neural underpinnings. Studies on statistical learning and behavioural stochasticity typically report the involvement of a widespread network of brain regions including sensory cortices and the cingulate cortex ^72,73^, as well as electrophysiological signatures with both early and late latencies ^23,74,75^. By contrast, sequences generated using artificial grammars or pattern repetitions and their violation typically recruit a more restricted set of brain regions including the inferior frontal gyrus and the hippocampus ^10,38,45,72,76,77^, and often elicit exclusively late electrophysiological signatures such as the P300 ^74,75^. The few studies that have directly contrasted both types of regularities, yet not fully orthogonally, confirm a dissociation in terms of anatomy ^27,39,41,47,48^ and timing of evoked responses ^74,75^.

Using different hypothesis spaces for regularity detection, as opposed to using a continuum, has important computational advantages for biological agents. Firstly, it allows for a faster detection of deterministic rules. This is because the deterministic rule hypothesis considers only extreme probability values (essentially 0 vs. 1), such that evidence for or against it accumulates rapidly ^28,57^. By contrast, the biases in higher-order transition probabilities that the *normative single-system model* tracks in order to detect rules take much longer to be detected.

Secondly, dividing the continuum of predictability into discrete hypotheses (as the *normative two-system model*) provides a tractable solution to the problem of regularity detection ^78,79^ and the representation of multifaceted hidden generative processes ^80^; a solution which seems favoured by humans. By contrast, computing with the continuum (as the *normative single-system model*) is computationally much more demanding, if not simply intractable. The distinct-system approach is currently explored in the field of general artificial intelligence ^81^ where algorithms combine local expert agents ^23,68,71,82^ that are specialized for different aspects of the learning process. This solution is much simpler and, when well-tuned, can prove much faster than the full optimal solution ^83,84^.

Thirdly, such a division of labor leverages dedicated computations for each hypothesis space, which can also reduce the total computational cost ^85^. In our case, the distinction between statistics and rules enables both a simple, accurate estimation of low-order statistical biases *and* a sensitivity to long-distance dependencies, with pattern matching. Tracking higher-order statistics, as in some versions of the *normative single-system model*, also preserves a sensitivity to long-distance dependencies but at the cost of a prohibitive growth of the number of transitions to monitor (they scale exponentially with the length of the dependency). By contrast, the number of patterns that are monitored at a given moment by the *normative two-system model* remains small. This is because all patterns that are incompatible with the sequence are excluded, resulting in at most one correct pattern per pattern length. Such a pruning strategy is very effective at reducing the cost of computations ^30,78,79,86^.

However, when using multiple hypothesis spaces, one must identify the one that is best suited for prediction or behavioural control, which requires arbitrating among hypotheses ^87–89^. We used a behavioural apparatus and instructions that advantageously left subjects unconstrained in the way they would arbitrate between the different hypotheses and found that they weighed these hypotheses in a graded manner: the graded comparison between (1) the random and statistical bias hypotheses was demonstrated by the specific detection dynamics of statistical biases and the false alarms, (2) the graded comparison between random and deterministic rule hypotheses by the specific detection dynamics of rules, (3) and the graded comparison between statistical biases and deterministic rule hypotheses by an interim competition before the detection of rules and by the graded weighing between them during random sequences.

These aspects of behaviour were better accounted for by the *normative two-system model*, than by an alternative *non-commensurable two-system model*, showing that subjects use a common probabilistic currency to arbitrate between the different hypotheses, in line with previous accounts of Bayesian inference applied to discrete states ^27,43,86^. By contrast, the *non-commensurable two-system model* computes pseudo posterior probabilities of the non-random hypotheses independently of each other and, hence, cannot normatively compare them. We showed that the *normative* and *non-commensurable models* make different predictions regarding categorisation profiles, detection dynamics of deterministic rules, and graded weighing of hypotheses, which all rejected *non-commensurable models*. It is worth noting that the only mathematical difference between the two models is the normalization term in Bayes’ rule which either includes all three hypotheses, or only two of them, respectively.

This normative graded weighing of hypotheses is all the more useful that both types of regularities can coexist in the same input, including speech ^16–18,90,91^. Deterministic rules can induce apparent statistical biases (e.g. the repetition of the AAB pattern induces globally more As than Bs), while statistical biases can induce local rule-like regularities (e.g. …ABABAB… in frequently alternating sequences). Many previous studies, by investigating statistics and rules separately, have neglected the possibility that one type of learning could interfere with the other, or foster regularity detection. Here, by manipulating the extent to which deterministic rules also induce apparent statistical biases, we found an interim competition between statistics and rules, even causing changes-of-mind ^57^, and characterized it (with Shannon entropy).

The normative graded weighing of hypotheses also explains sequence miscategorisations and other aspects of behaviour which appear suboptimal at first sight. For instance, subjects misjudged sequences with a statistical bias as fully random when the evidence, as inferred by the *normative two-system model*, was indeed weaker. Furthermore, subjects failed to detect alternation biases more often than repetition biases, and biases in item frequency, which is in line with past studies showing that the sequences that humans judge as maximally random in fact alternate more than expected by chance ^54,61^. Also, during the course of the sequence, subjects’ false alarms were explained by local, mostly statistical, departures from randomness detected by the *normative two-system model*, as previously observed ^67^. In addition, past studies reported that humans are prone to seeing regularity when there are actually none ^54,61–63^. Accordingly, we also found that subjects often categorized fully random sequences as entailing a statistical bias. The *normative two-system model* also showed such miscategorisations, yet to a smaller extent. The larger effect found in subjects (also for the false alarms in the course of a sequence) may come from a general underweighting of large probabilities (in favour of the random hypothesis) and an overweighting of small probabilities (in favour of the non-random hypotheses), a bias vastly reported by Kahneman & Tversky ^92^ which we also observed here (*Supplementary Note 1*); or, alternatively, a non-uniform prior over random vs. non-random hypotheses.

We now acknowledge several limitations of our study. A first limitation is that we used a restricted set of regularities, due to experimental time constraints. This restricted set enabled us to study the estimation of first-order transition probabilities and detection of repeating patterns, but is not suited to study more complex types of regularity ^2^ that require more complex computations ^46^. Notably, humans can detect deterministic rules such as A^*n*^B^*n*^ with an increasing number *n* (e.g. AB-AABB-AAABBB…). A possibility, which has received some behavioural support, is that the human brain uses a ‘language of thought’ combining logical, numerical and geometrical primitives in order to compress deterministic sequences into a regular expression ^42,44,93–95^. For simplicity, we manipulated here only repeated patterns of various lengths, but found, in line with a possible role of compression, that the barely compressible length-10 pattern AAABAABBAB was more often missed by subjects compared to the other two length-10 patterns, AAAAABBBBB and to a lesser extent, BABABABBAA, which are more compressible ([A^4^, B^4^] and [A^2^, (BA)^2^, B^2^] respectively). Note, however, that this effect could also result from the difference in the patterns’ apparent statistical bias (which is weak in AAABAABBAB and strong in AAAAABBBBB), an alternative we have recently considered and discarded ^95^.

This brings us to the more general question of what is psychologically general in the *normative two-system model* and what is specific to the task. Although we can only speculate about the scope of statistical biases (i.e. which order(s) of transitions) and deterministic rules (i.e. which type(s) of rules, beyond repetition) which is relevant for the brain, the model makes the general claim that there exist two fundamentally distinct categories of regularity, statistics and rules, which afford respectively uncertain vs. sure predictions. The model also posits that the statistical bias and deterministic rule hypotheses are evaluated in parallel during sequence processing, leading to testable predictions ^95^. The model finally posits that humans can rationally compare those different hypotheses using a common probabilistic currency. Future studies should thus investigate a more diverse and ecological set of regularities and inspect whether humans conform to the model’s predictions also when the variety of regularities is much wider but could still fall into our proposed taxonomy.

## Methods

### Sequences’ generative process

Each sequence was composed of 200 binary observations, that we refer to as A or B, and generated by one of the following 3 generative processes: (1) they could be random from the beginning to the end, (2) with a statistical bias introduced after an initial random part, (3) with a deterministic rule introduced after an initial random part.

#### Random parts

The random parts of sequences were generated by independent draws with equal probability for either binary outcome (i.e. as tosses of a fair coin). The resulting sequences contain on average as many As as Bs and as many repetitions (i.e. AA and BB) as alternations (i.e. AB and BA).

#### Change-points

In sequences entailing a regularity, a change-point separates the initial, random, part from the second, non-random, part. Unbeknownst to the subjects, the position of the change-point was drawn from a gaussian distribution centred on the middle of the sequence (i.e. observation #100) with a standard deviation of 15 observations, truncated from observations #55 to #145 such that the change would not appear neither too early nor too late in the sequence. The resulting empirical distribution of change-points is centred on observation #100.65 ± 15.55 SD.

#### Statistical biases

Statistical regularities are obtained by drawing observations from a biased first-order Markov chain. Each statistical bias is thus fully described by the values of two first-order transition probabilities: *p*(A|B) = 1 − *p*(B|B) and *p*(B|A) = 1 − *p*(A|A). Note that unless otherwise specified, all transition probabilities are first-order. Importantly, transition probabilities also fully determine lower-order statistics: the frequency of items, *p*(A) = 1 − *p*(B), and the frequency of alternation, *p*(alternation) = 1 − *p*(repetition) ^23,54^. We used the following statistical biases (the numbers indicate (*p*(A|B), *p*(B|A))): more repetitions than alternations with (1/3, 1/3), (1/4, 1/4), and (1/5, 1/5); more alternations than repetitions with (2/3, 2/3), (3/4, 3/4) and (4/5, 4/5); more of one item than the other with (1/3, 2/3), (1/4, 1/2) and (1/5, 4/5), which also bias the alternation frequency; or biased both in terms of alternation and item frequencies with (1/2, 1/4) and (3/4, 1/2). For repetition biases and alternation biases, the corresponding frequency of items is at chance level. We quantified the strength of the statistical biases as the Shannon entropy (i.e. *H*) of the distribution of pairs of items ^96^

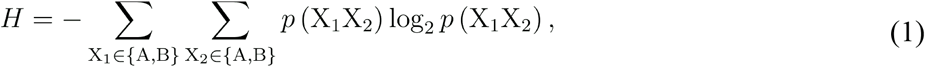

where the subscripts denote the position of the observation within the pair. The probability of a given pair of items can be computed using the generative transition probabilities

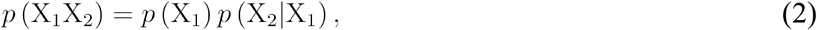

with

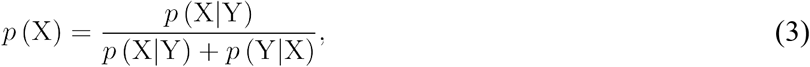

where X is either A or B and Y is either B or A. The entropy of the chosen statistical biases ranged from *H* = 1.44 to 1.92 bit, which contrasts with the entropy characterising random sequences, which is maximal, that is *H* = 2 bits. We used the limit 0 × log_2_(0) = 0 when it applied.

#### Deterministic rules

Deterministic rules were obtained by repeating a fixed pattern. Different pattern lengths were used: 4, 6, 8 and 10 observations. Moreover, patterns were chosen depending on the strength and type of bias in the apparent transition probabilities they induce. We term this bias ‘apparent’, in opposition to ‘generative’ since the deterministic rules are not generated according to a statistical bias, but indeed by repeating a pattern. Nonetheless, depending on the pattern used, a statistical bias may emerge: no bias with AABB and AAABBABB; a repetition bias with AAABBB, AAAABBBB and AAAAABBBBB; an alternation bias with AABABB, AABABABB and AABABABABB; and a bias in the frequency of items with AAABAB and AAABAABBAB. We selected several of these patterns, both in terms of bias type and bias strength so as to match with most of the biases characterising the statistical regularities we used. Apparent transition probabilities can be computed for each deterministic rule (supposing an infinite number of repetitions of that pattern) and then used to estimate Shannon entropy, quantifying the strength of their apparent statistical bias using *Equation 1*.

### Experimental protocol

#### Subjects

A total of 28 subjects participated in the study. Data from 5 subjects were rejected because of very poor data quality reflecting a lack of comprehension of the instructions. Note, however, that our conclusions hold when using data from all the subjects; the reader can rerun our analyses with the code we provide online together with the full dataset. The data presented here come from 23 subjects (16 females) aged between 20 and 29 years old (mean age 23.91 ± 2.59 SD) from various education backgrounds including history of arts, law, translation, biology, engineering, psychology, etc. All subjects were right-handed. The study was approved by an independent ethics committee (CPP 08-021 Ile-de-France VII), and subjects gave their informed written consent before participating.

#### Stimulation

Stimuli consisted of two tones composed of three superimposed sine waves (350, 700, and 1400 Hz vs. 500, 1000, and 2000 Hz). The tones were 50 ms long, with 7 ms rise and fall times. They were randomly associated to A or B before each sequence. The stimulus-onset asynchrony was 300 ms long, thus resulting in 3.33 sounds per second. Stimulation was delivered using MATLAB and Psychtoolbox ^97^.

#### Triangular arena

The triangular arena is an equilateral triangle displayed on a touch screen and whose vertices correspond to the 3 possible generative hypotheses: the bottom vertex corresponds to a random process and the two upper vertices correspond to non-random processes (i.e. statistics and rules) whose respective side (i.e. left or right) was counterbalanced across subjects. For the sake of averaging and clarity of reporting, the trajectories of half of the subjects were thus vertically mirrored such that for all subjects the statistical bias hypothesis corresponds to the left upper vertex and the deterministic rule hypothesis to the right upper vertex. A point in that triangular space can be converted into posterior probabilities for each generative hypothesis (by means of barycentric coordinates summing to 1, see below). The centre of the triangle corresponds to equal probabilities (i.e. (1/3, 1/3, 1/3)). Because all sequences start with a random part, observers started at the bottom vertex and updated their locations in the triangle as observations were delivered in a way that reflects their estimates of the probabilities for the three possible hypotheses given the received observations. Subjects were allowed to move freely inside the triangular arena: they could pause at a particular location of the triangle or change their mind for instance by returning near the random vertex after a suspected regularity was discarded. The triangular arena (9.7 cm wide × 8.4 cm high, 800 pixels wide × 693 pixels high) was displayed using MATLAB and Psychtoolbox ^97^ on a 14 inches touch screen (31 cm wide × 17.4 cm high, 2560 pixels wide × 1440 pixels high) computer (HP Pavilion x360) that was lying horizontally on a table. Finger position was collected using MATLAB immediately after each auditory stimulus was played, thereby resulting in a trajectory composed of 200 pairs of cartesian coordinates (*a_k_* and *b_k_*). When the recorded finger position was outside of the triangle (because of subject’s motor imprecision), the positions were projected back to the nearest point on the triangle border (through Euclidean distance minimization, average distance = 5.83 pixels ±3.88 SD, [1.77, 16.0]).

#### Training

Subjects were first given written instructions. They were then presented with several short example sequences (70 observations, no change-point) and told the underlying generative process in order for them to understand the differences between the two types of regularities and the random process. The experimenter provided examples of how to use the triangular arena to report probabilistic beliefs with 4 example sequences (1 random, 2 random-to-statistical and 1 random-to-rule). Finally, subjects practiced on 7 example sequences (2 random, 3 random-to-statistical and 2 random-to-rule). The experimenter then answered possible questions and launched the experiment. The regularities used during training were different from those used in the experiment.

#### Experiment

During the experiment, subjects were presented with 33 sequences (10 random, 13 random-to-statistical and 10 random-to-rule) whose order was randomly defined for each subject. These sequences were allocated to 4 different sessions (each composed of 8 or 9 sequences) interspersed with pauses. Two of the random-to-statistical sequences were duplicates (of (1/2, 1/4) and (3/4, 1/2)) that were added in order to assess within-subject variability in stochastic conditions. They are simply discarded from the analyses presented here. All trials follow the same structure. Subjects initiated the trial by touching a button on the touch screen. The triangular arena was then displayed on the screen and a 3-second countdown was initiated. During the following minute, the sequence was played, and subjects moved their finger inside the triangular arena. Once the presentation of the sequence finished, subjects were asked several post-sequence questions. They were first asked whether they thought retrospectively that the sequence entailed a regularity or not (detection). In the absence of noticed regularities, the trial was terminated. Otherwise, they were asked several other questions. Firstly, which type of regularity (statistics vs. rule) occurred (discrimination)? Secondly, when the change-point occurred? In that case, they used a continuous horizontal scale from observation #1 to #200 and also rated their confidence in that estimate using a continuous vertical scale. Those two questions are in retrospect, which is in contrast with the third question in which they were asked to estimate when they had realized, during the sequence presentation, that a change-point had occurred. In that case, they also used a continuous horizontal scale from observation reported as change-point (at the second question) to observation #200 and also rated their confidence in that estimate using a continuous vertical scale. The data from the third question were not used here.

### Normative two-system model

We derive the ideal observer model for the task, which is the Bayes-optimal solution using the actual task structure ^98,99^. Comparing subjects behaviour against this benchmark inference therefore allows the identification of signatures of a normative inference in human behaviour ^59,60^. The model is presented with a given sequence *y*, of length |*y*| = *K*, and returns the posterior probability of each possible hypothesis *H_i_* (i.e. random, random-to-statistical and random-to-rule processes) using Bayes’ theorem, as the sequence unfolds in time

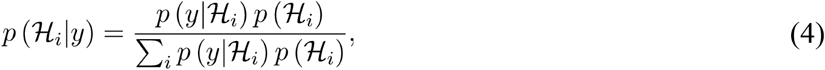

The assumptions of the model closely correspond to the task structure, and they determine its prior and likelihood functions. The likelihood of the sequence under each hypothesis *p*(*y*|*H_i_*) is derived below. The prior probability over hypotheses was uniform, such that *p*(*H_i_*) = ⅓.

#### Sequence likelihood under a random hypothesis

Under the random hypothesis *H*_rand_, all observations are at chance level and independent. The likelihood of a sequence is thus the product of the chance-level likelihoods of each observation. Because the observations are binary, the process amounts to independent tosses of a fair coin, and the likelihood of each observation is ½,

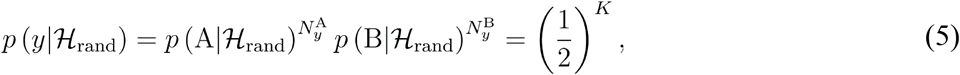

where *N_y_*^X^ denotes the number of times X has been observed in the sequence *y*.

#### Likelihood of a sequence under non-random hypotheses

For the remaining two hypotheses, a change-point delineates the initial, random, part of the sequence from the second, non-random, part. However, the location of the change-point is unknown and assumed random. In order to get rid of this unknown factor, one must therefore consider all possible *j_k_* positions of the change-point (i.e. after the 1^st^, 2^nd^, …, up to the (*K* − 1)^th^ observation), and marginalize over these positions

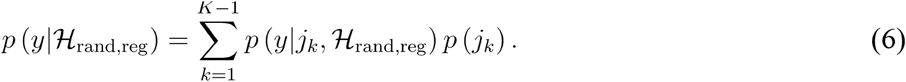

The likelihood of the sequence is, for a given change-point location, the product of the likelihood of the random part and the non-random part of the sequence

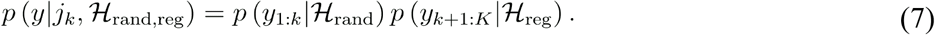

The likelihood of the first part of the sequence (*y*_1:*k*_, noted below *y*_rand_ for brevity) is the likelihood of a random sequence (*Equation 5*), the likelihood of the second part (*y*_*k*+1:*K*_, noted below *y*_stat_ or *y*_rule_ for brevity, whose length is |*y*_stat or rule_| =*K* − *j_k_*) depends upon the type of regularity that is considered (either statistical bias or deterministic rule, see below). The prior distribution over change-point positions is set to be uniform, such that all positions are a priori equally likely *p*(*j_k_* = 1/200.

#### Sequence likelihood under a statistical bias hypothesis

The random-to-statistical hypothesis *p*(*y*|*H*_rand,stat_) considers that, after the change-point, the sequence likelihood depends on a matrix of transition probabilities between successive observations characterising a first-order Markov chain ^54^. The generative matrix of transition probabilities is unknown and random from one sequence to the next. In order to get rid of this unknown factor, one must therefore consider all possible values of transition probabilities ***θ*** and marginalize over them. The likelihood of the second, statistical, part of the sequence *y*_stat_ is thus

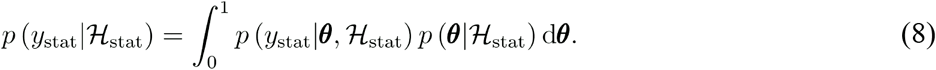

For a model that estimates transition probabilities between consecutive stimuli, the likelihood of a given observation depends only on the estimated transition probabilities and the previous stimulus. For simplicity, the first observation can be considered as arbitrary such that its likelihood equals chance level. This results in

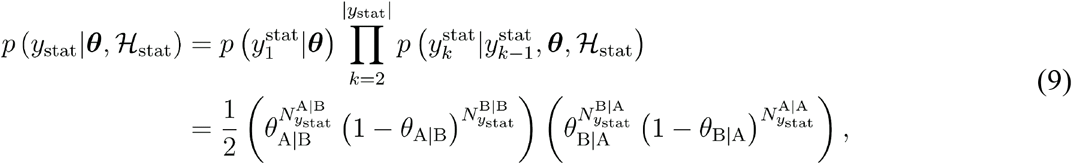

where ***θ*** is a vector of two transition probabilities ***θ*** = [*θ*_A|B_, *θ*_B|A_] and *N*^X|Y^ denotes the number of YX pairs in the statistical part of the sequence. *Equation 9* corresponds to the product of two Beta distributions, which parameters are the transition counts plus one. Inserting *Equation 9* in *8*, and assuming a flat prior, we obtain

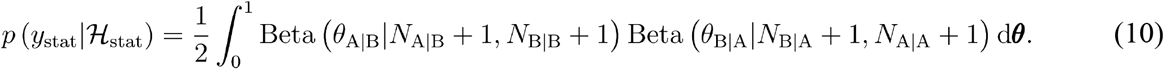

The integral over such Beta distributions has analytical solutions that can be used to compute sequence likelihood in *Equation 8* when a uniform conjugate prior over ***θ*** is used ^100^.

#### Sequence likelihood under a deterministic rule hypothesis

The random-to-rule hypothesis *p*(*y*|*H*_rand,rule_) considers that, after the change-point, the sequence can be described as the repetition of a particular pattern. However, this pattern is unknown. In order to get rid of this unknown factor, one must therefore consider all possible patterns, denoted {*R*}, and marginalize over them. The likelihood of the second, deterministic, part of the sequences *y*_rule_ is thus

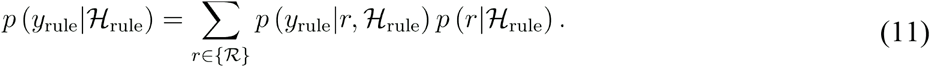

We consider that the set of all patterns {*R*} can be split in two disjoint subsets. The first one corresponds to all patterns whose length is equal or smaller than the length of the deterministic sequence: this is the set of patterns potentially entirely observed in the current sequence {*R*^o^}. The second subset corresponds to patterns whose length is larger than the length of the deterministic sequence: this is the set of partially observed patterns {*R*^u^}. The size of that latter set depends upon the maximum pattern length allowed, denoted *l*. We used *l* = 10 observations such that the model can detect all the patterns used in the experiment while remaining psychologically plausible given human memory limits. This parameter choice actually does not significantly impact the predictions of the model for *l* > 10 as it allows the detection of all patterns used here (*Extended Data Fig. 8*). For both patterns’ sets, the likelihood of a particular pattern *r* given the observed sequence is all or none

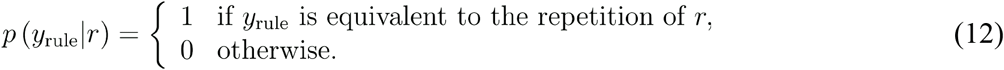

Among {*R*^o^}, there is at most one (and thus possibility zero) pattern of each given length that is compatible with the observed sequence (i.e. for which the sequence likelihood is not null). Possible extensions of this pattern found in {*R*^u^} also result in a sequence likelihood equal to one, and all the others patterns are incompatible with the observed sequence (and thus result in a likelihood that is null).

The prior probability of patterns reflects an incremental procedure for pattern generation: at each iteration, the current pattern is either selected, which terminates the procedure, or extended with observation A or B, with equal probability among those 3 possibilities. The process terminates when the maximum pattern length *l* is reached. With such a process, longer patterns are less probable than shorter ones, the prior therefore favours shorter patterns (see first ratio in *Equation 13*), following the ‘size-principle’ which has been widely used in the past ^101^. For a given pattern *r* of length |*r*|, the prior probability is

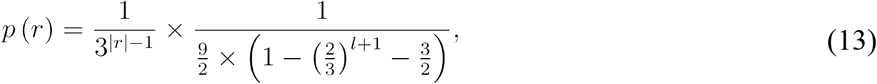

where the second ratio ensures, given that *l* is the maximum pattern length allowed, that ∑_*r*∈{*R*}_*p*(*r*) = 1.

#### Posterior distribution of change-point position

Besides computing the posterior probability of the three different generative processes given an input sequence, it is also possible to compute the posterior probability of the change-point being at one particular position, for the two hypotheses that assume the existence of such a change-point, that is

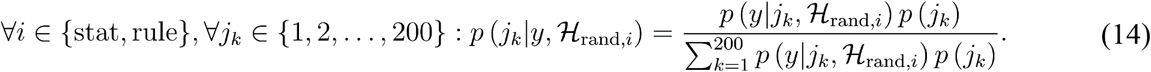

The likelihood of an observed sequence containing a change-point has been previously defined in *Equation 7*. By contrast, if the change-point is in the yet unobserved part of the sequence, then the current sequence is still necessarily random:

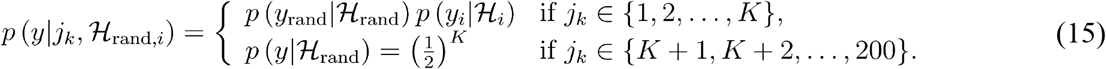

Those hypothesis-specific posterior distributions over change-points can then be combined by means of Bayesian model averaging

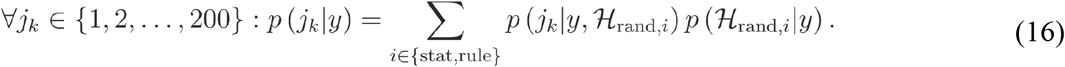

The most likely position of the change-point is therefore the maximum a posteriori of this distribution and the confidence related to that estimate is assessed by measuring the log-precision of the posterior distribution ^24,55,56^.

### Normative single-system model

An alternative to the dichotomy between the statistical bias hypothesis (with order 1 transition probabilities) and the deterministic rule hypothesis (with patterns) is that subjects monitor transition probabilities in all cases (*H*_rule_ and *H*_stat_) and use apparent biases in (possibly higher-order) transition probabilities that characterize deterministic rules to detect them. We explored transition probabilities of various orders, also referred to as *n*-grams models in the linguistic literature ^102^. We also allowed non-uniform (conjugate) priors that promote predictability in the transition probabilities, by introducing a parameter *d*: *d* = 0 corresponds to a uniform prior (the next observation is expected with any probability) while *d* > 0 favours predictability (the next observations is expected with a probability closer to 0 or 1) ^103^. The likelihood of the non-random part of the sequence becomes

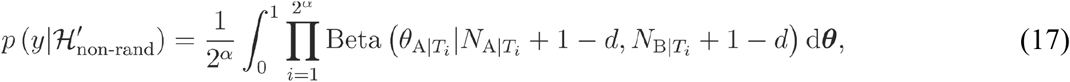

where *α* is the order of the chain, resulting in 2^*α*^ possible transitions **T**. For instance, a Markov chain of order *α* = 1 corresponds to the transitions **T** = [X|A, X|B], a chain of order *α* = 2 considers the transitions **T** = [X|AA, X|AB, X|BA, X|BB], etc. We present results only for *d* = 0.001, because larger values quickly make the *normative single-system model* worse than the *normative two-system model*. The likelihood function we used for the statistical bias hypothesis (*H*_stat_) of the *normative two-system model* (*Equation 10*) is thus a spatial case of *Equation 17* where *α* = 1 and *d* = 0, that is a first-order Markov chain with a uniform prior.

We adopted a factorial approach to evaluate all possible variants of the *normative single-system model:* on the one hand, *H’*_stat_ and *H’*_rule_ could both correspond to the same transition order 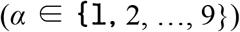, or to different transition orders (*α*_stat_ = 1, and 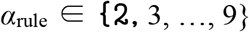); on the other hand, *H’*_stat_ and *H’*_rule_ could both use the same prior belief about predictability (*d* = 0), or different priors (*d*_stat_ = 0, and *d*_rule_ > 0). Combining those two dimensions (order and prior) gave rise to four families of models, and for each of which we varied *α* (and *d* when relevant). In three of those four families, the comparison of hypotheses remains normative (as in *Equation 4*). When *H’*_stat_ and *H’*_rule_ resort to identical transition order and prior belief, they become strictly identical, preventing normative categorisation in the task. This family of models must thus rely on a heuristic in order to compare between non-random hypotheses. For this, we used the distance between the prediction of the non-random hypothesis and chance: |½ − *p*(*y*_*K*+1_|*y*_1:*K*_, *H*_non-rand_)|, where

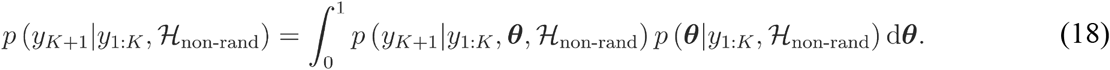

This distance is then used to weigh *H’*_stat_ and *H’*_rule_ (tying *H’*_rule_ to smaller distances, corresponding to increased predictability) either linearly or not (*Extended Data Fig. 6*).

### Non-commensurable two-system model

The normative arbitration between the non-random hypotheses requires a common probabilistic currency. Another alternative model is thus that subjects evaluate the posterior probability of each non-random hypothesis (against the random hypothesis) independently of one another, making them non-commensurable. To explore this possibility, and contrary to the *normative two-system model*, we computed posterior probability of each non-random hypothesis separately, given the observed sequence

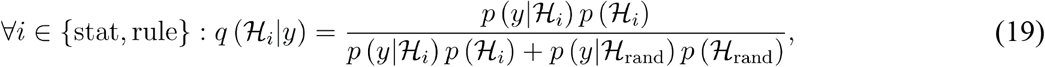

where the prior probabilities are set to chance level (i.e. ½ in that case). From *Equation 19*, it follows that the probability of the random hypothesis depends upon the non-random hypothesis that is considered

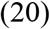

In order to report the probabilities of the 3 hypotheses in the triangular arena (which requires they sum to 1), we used four different combination functions: (1) relative weight, (2) selection of maximum (taking the best hypothesis and discarding the other), (3) a sigmoid based on the difference between the posterior probability of non-random hypotheses, and (4) a sigmoid based on the log-ratio between the posterior probability of non-random hypotheses. For the two latter cases, one parameter (*β*) controls the slope of the sigmoid function (we explored 100 values equally spaced logarithmically between 1 and 10^8^).

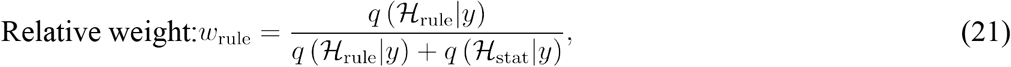

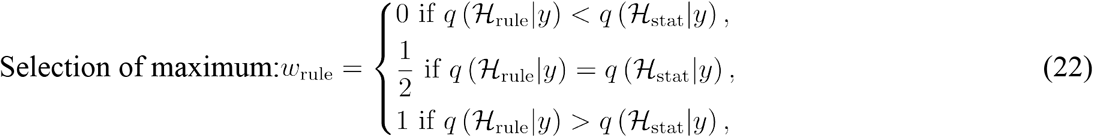

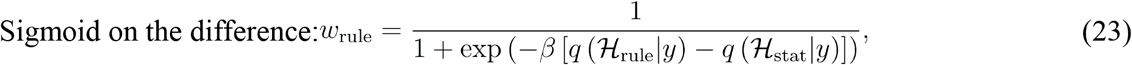

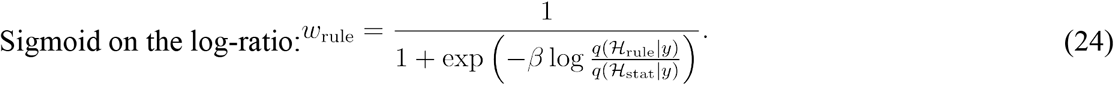

The pseudo posterior probability of the random hypothesis is first obtained by a weighted combination of the probability of the random hypothesis under each non-random hypothesis

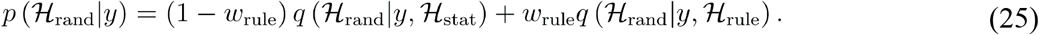

Then, the pseudo posterior probabilities of the non-random hypotheses are computed based on the weights from the previously mentioned combination functions which guarantees that the probabilities over the 3 hypotheses sum to 1 as

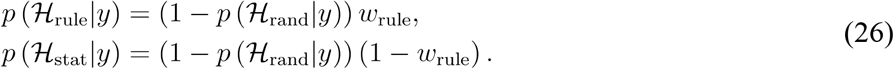

### Sequences included in the analyses

Analyses of categorisation profiles were performed on all sequences. All other analyses of non-random sequences were restricted to sequences with regularities that were accurately detected so as to ensure that the effects we report were not due to a miscategorisation of sequences. Analyses inspecting dynamics locked on detection-points (i.e. when the correct hypothesis becomes more likely than the other two together) were further restricted to non-random sequences for which such a detection-point could be found. Analyses of random sequences were performed on both the initial part of random-to-non-random sequences, and the entire length of random sequences.

### Signal detection theory

The detection sensitivity index *d’*_detect_ ^104^ between random and non-random sequences is computed by comparing non-random sequences accurately labelled as non-random vs. random sequences erroneously reported as non-random using answers to the first post-sequence question (presence of regularity?)

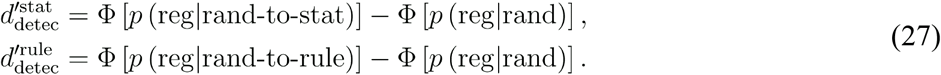

where Φ is the inverse cumulative function of the gaussian distribution (with *μ* = 0 and *σ* = 1). By contrast, the discrimination sensitivity index *d’*_discr_ ^104^ between the two random-to-non-random sequences is computed by comparing correct to incorrect categorisations of regularities at the second post-sequence question (type of regularity?)

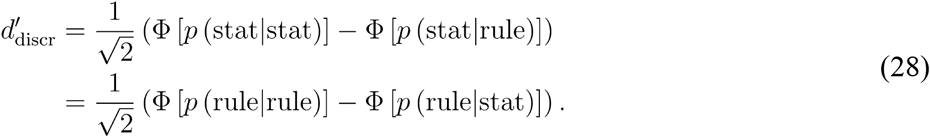

### From finger position to posterior probability

A given pair of cartesian coordinates (*a, b*) specifying the subject’s finger position can be turned into the posterior probability of each hypothesis *p*(*H_i_*|*y*) by performing a conversion to the barycentric coordinate system using the following equations:

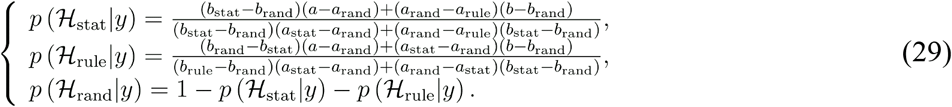

Where *a_h_* is the horizontal position of hypothesis *h* on the screen and *b_h_* its vertical position (both in pixels). Conversely, the posterior probability of the three hypotheses estimated by the model can be reported on the touch screen by applying a barycentric-to-cartesian transformation

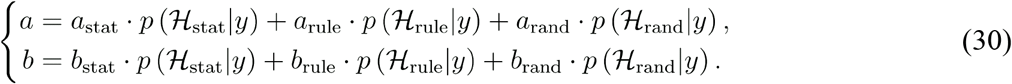

### Triangular histograms of positions in the triangle

The histograms were obtained by segmenting the triangle into 87 vertical and 101 horizontal rectangular parcels (since the height of an equilateral triangle is 0.87 times the size of its edge) and counting the frequency of occupancy of each of these parcels. Before counting, the subjects’ and model’s trajectories were linearly interpolated (resulting in 3 times more data points). The resulting 2D histogram was then smoothed with a gaussian kernel (*σ* = 4) and normalized such that the sum over parcels equals 1.

### Measures of dynamics

We used different measures to quantify different aspects of the inference dynamics.

#### Detection-point

Detection-points are defined as the number of observations after the change-point for the posterior probability in the correct hypothesis to become more likely than any of the others (i.e. larger than ½).

#### Detection slope

A sigmoid function was used to approximate the slope of detection dynamics

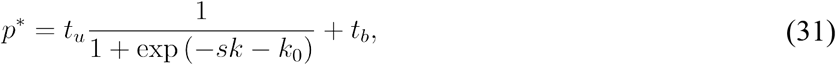

where *s*, *k*_0_, *t_u_* and *t_b_* are the 4 free parameters of the function (respectively the slope, the intercept and the upper and lower bounds) that transforms the position *k* of an observation in the sequence into a given probability estimate *p*^*^. They were fitted using a grid search technique that finds the set of 4 parameters that minimizes the MSE along the following parameter grid: for the slope parameter, 31 values logarithmically equally spaced between decades 10^−3^ and 10^0^; for the intercept parameter, any observation number between 0 (change-point) and 200; for the offset parameter, 11 values linearly spaced between 0 and 0.5 with a step of 0.05; for the scaling parameter, 11 value linearly spaced between 0.5 and 1 with a step of 0.05.

#### Belief update

The amount of belief update quantifies the average increase in posterior probability each observation in the non-random part of the sequence provided to the correct non-random hypothesis

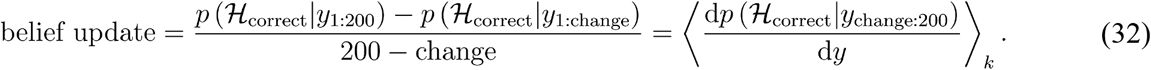

#### Belief difference

An index reflecting the competition among the non-random hypotheses (statistical bias vs. deterministic rule) independently from the posterior probability of the random hypothesis was also computed

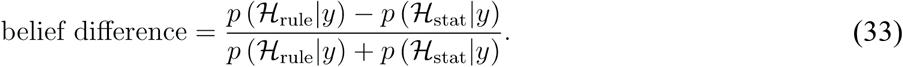

This index ranges from −1 (statistical bias) to 1 (deterministic rule). When both non-random posterior probabilities were null, we set belief difference to 0.

### Model comparison

To arbitrate between different models of sequence processing, we measured the mean squared error between specific features of subjects’ behaviour and the corresponding feature in models. Measuring the error coarsely on the full time-series of posterior probabilities reported by subjects and models in all sequences would have led to conclusions mostly driven by the rate of temporary false alarms (because most observations are generated purely at random) rather than by the detection of regularities. We thus focused on several specific features (detailed below) which we found most informative in order to distinguish between models.

#### Categorisation profile

We quantified the error in sequence categorisation. Models identify the most likely generative process based on the posterior probability of each hypothesis at the end of the sequence. When models were undecided (i.e. when two hypotheses have probability ½ ± 0.001 or the three hypotheses have probability ⅓ ± 0.001; the findings are robust to the precision parameter), models chose randomly among the most likely hypotheses.

#### Smoothness of deterministic detection

We quantified the smoothness versus unsteadiness of the detection dynamics for deterministic rules by averaging the absolute value of the second-order derivative of the posterior probability of the deterministic rule hypothesis (average over all observations after the change-points). Stronger competition between non-random hypotheses during the detection of deterministic rules exhibiting an apparent statistical bias result in frequent acceleration-deceleration in the *single-system model*, and thus high average absolute acceleration.

#### Fit of belief difference

We quantified the error of a linear regression relating the model’s belief difference to the subjects’ (after regressing-out the effect of confounding variables such as the posterior probability of the random hypothesis, and together with its interaction with the posterior difference in the regression). We restricted the fit to null or positive regression coefficients (a negative value would denote opposite ranking of *H*_rule_ and *H*_stat_ between models and subjects).

### Statistics

All dispersion metrics reported in the figures are standard errors of the mean (s.e.m., with *n* = 23 subjects), except shaded areas around regression lines which reflect 95% confidence intervals around estimated regression coefficients. Reported correlation coefficients (noted *r*) are Pearson coefficients. Parametric frequentist statistics with repeated measures are performed. For the *t*-tests, the mean value of the paired difference, 95% confidence intervals, an estimate of effect size (Cohen’s *d*), the *t*-statistic, the corresponding degrees of freedom (noted *t*_df_) and *p*-values (noted *p*) are reported. For the ANOVAs, an estimate of effect size (*w*^2^), the value of the *F* statistic, the corresponding degrees of freedom (noted *F*_df_ and *p*-values (noted *p*) are reported. All statistical tests were two-tailed at with a type I error risk of 0.05. Bayesian *t*-tests were used to quantify the evidence in favour of the null hypothesis using a scale factor of 0.707 ^105^.

## Code availability

The MATLAB code used to run simulations of the different models, analyse the results and reproduce all the figures is available on *GitHub* (https://github.com/maheump/Emergence).

## Data availability

The dataset presented in the current study is available on *GitHub* (https://github.com/maheump/Emergence).

## Acknowledgements

M.M. was supported by a ‘*Frontières du Vivant’* doctoral fellowship involving the *Ministère de l’Enseignement Supérieur et de la Recherche* and the *Fondation Bettencourt Schueller*, and a *Fondation pour la Recherche Médicale* doctoral fellowship. This research was funded by *Institut National de la Santé Et de la Recherche Médicale* (to S.D.), *Commissariat à l’Energie Atomique* (to S.D. and F.M.), *Collège de France* (to S.D.), a *European Research Council* grant *‘NeuroSyntax’* (to S.D.).

We would like to thank the subjects that participated in the study as well as Isabelle Brunet for her help in data acquisition. We thank Lucie Berkovitch and Jacques Pesnot-Lerousseau for help in piloting the experiment. We are also grateful to Athena Akrami and her group, Maria Chait, Philippe Domenech, Stephen Fleming and his group, Luc Mallet and his group, Karim N’Diaye, Emmanuel Procyk, Jérôme Sackur, Mariano Sigman, Valentin Wyart, and the members of the *Cognitive Neuroimaging Unit* (*NeuroSpin*) for useful discussions.

## Author contributions

Conceptualization: M.M., F.M., S.D.; Formal analysis: M.M., F.M.; Funding acquisition: S.D.; Investigation: M.M.; Methodology: M.M.; Project administration: M.M.; Software: M.M.; Supervision: F.M., S.D.; Visualization: M.M.; Writing — original draft: M.M.; Writing — review & editing: M.M., F.M., S.D.

## Additional information

**Extended data** is available for this paper.

**Supplementary information** is available for this paper.

**Competing interests:** The authors declare no competing interests.

**Correspondence** should be addressed to M.M.

## Extended data figures

**Extended Data Fig. 1.**
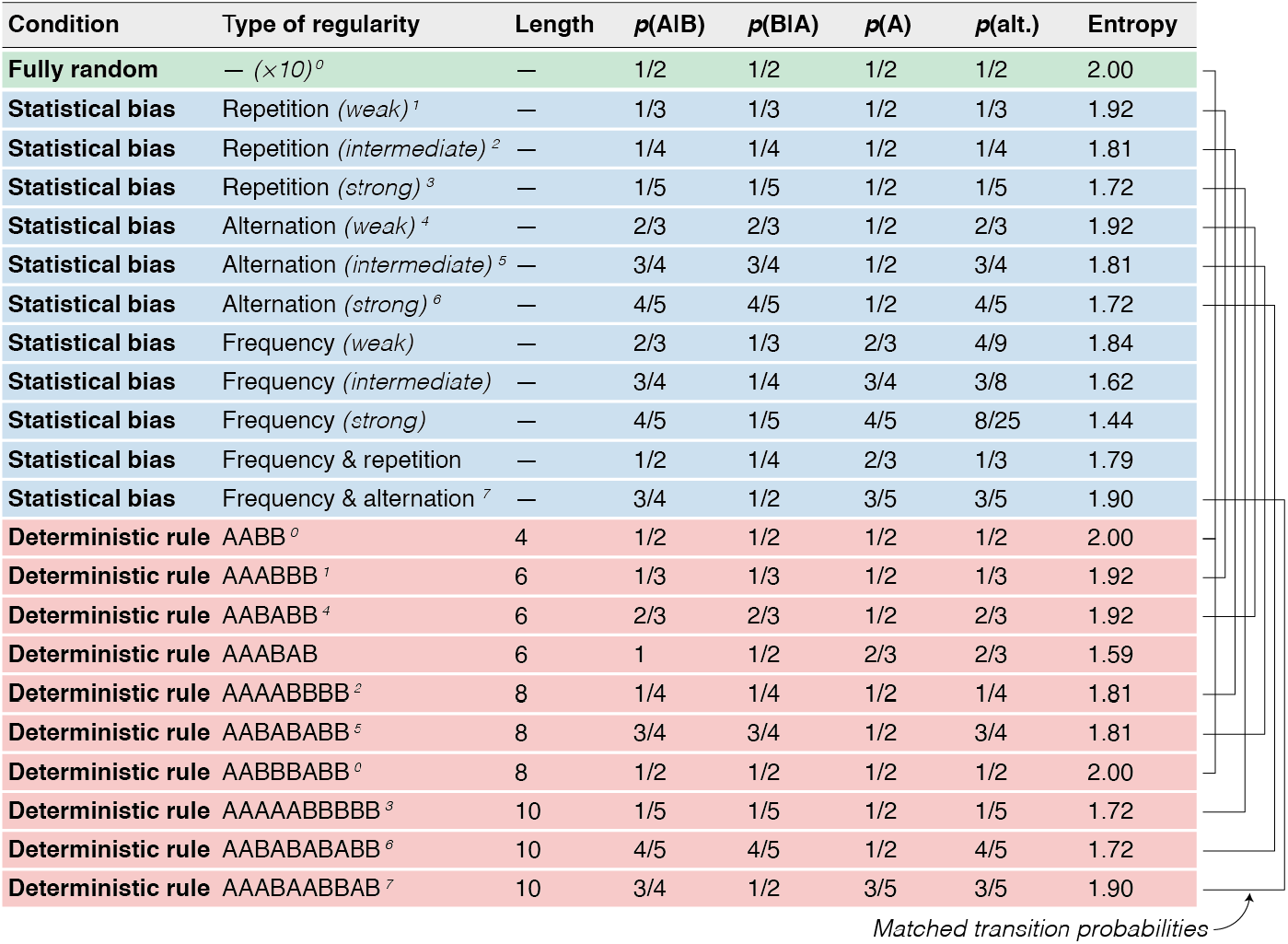
Experimental conditions. *H* corresponds to the Shannon entropy of generative transition probabilities (in bits). In the case of statistical biases, the type of bias (i.e. repetition-, alternation- and/or frequency bias) and its strength (i.e. weak, intermediate or strong) is specified. Superscripts indicate sequences with matched (apparent) transition probabilities, also illustrated by the tree on the right border.

**Extended Data Fig. 2.**
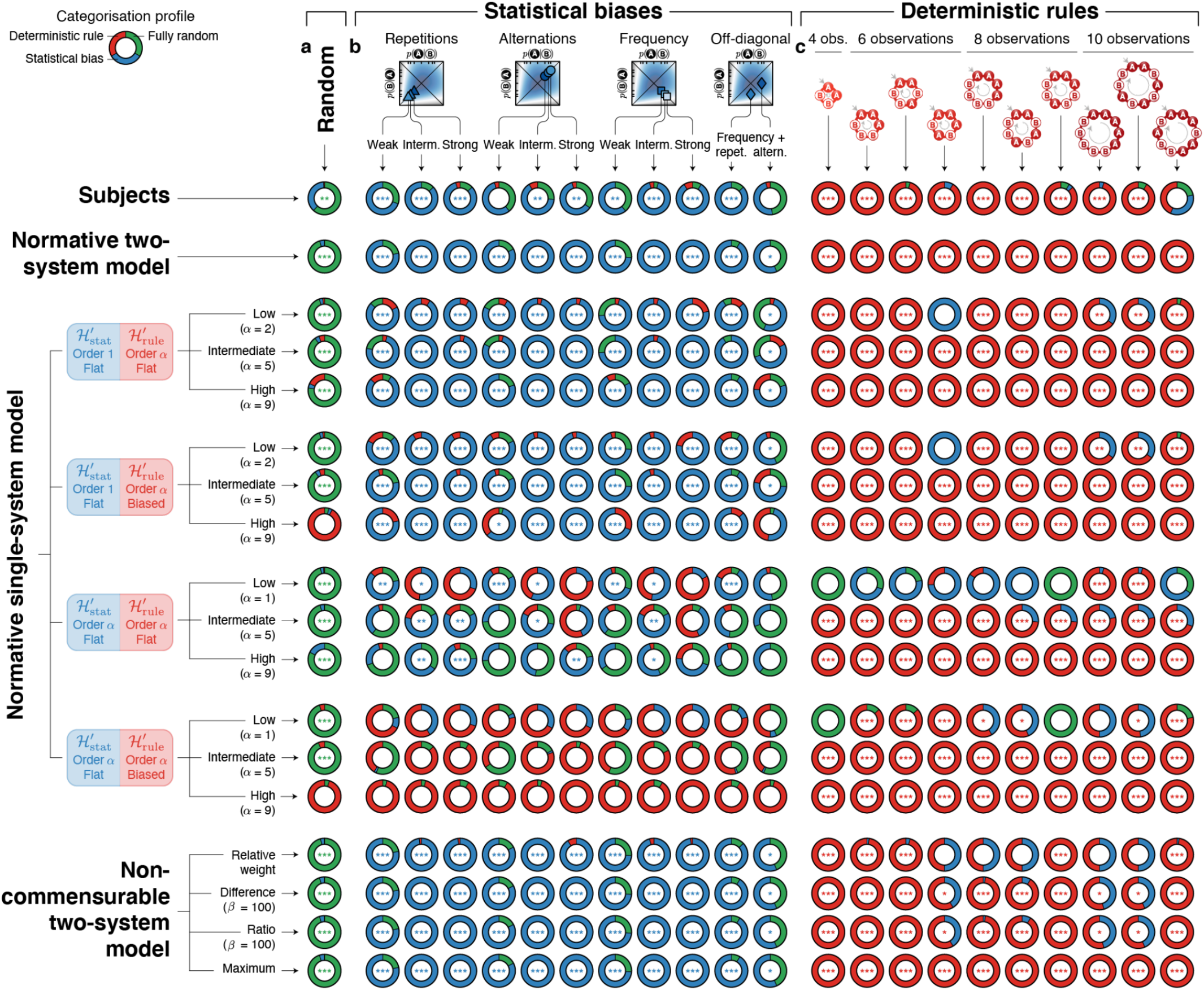
Categorisation of sequences by subjects and models. Proportion of generative processes reported by subjects and the different models for each sequence. In the case of fully random sequences (**a**), sequences with statistical biases (**b**), or a deterministic rule (**c**), responses are averaged over sequences (*N*_seq_ = 10). Subjects report the generative process using post-sequence questions. Models identify the generative process based on the maximum a posteriori probability over hypotheses at the end of the sequence. When the model is undecided (e.g. when *p*(*H*_rule_|*y*) ≈ *p*(*H*_stat_|*y*) ≈ ½; with a precision of 0.001), the model chooses randomly among hypotheses. *H* corresponds to the Shannon entropy of generative transition probabilities (in bits). The probabilities *p*(A) and *p*(alternation) are analytically computed from generative transition probabilities. Stars denote significance of an exact binomial one-tailed test of the proportion of correct categorisation larger than ⅓: *** *p* < 0.005, ** *p* < 0.01, * *p* < 0.05.

**Extended Data Fig. 3.**
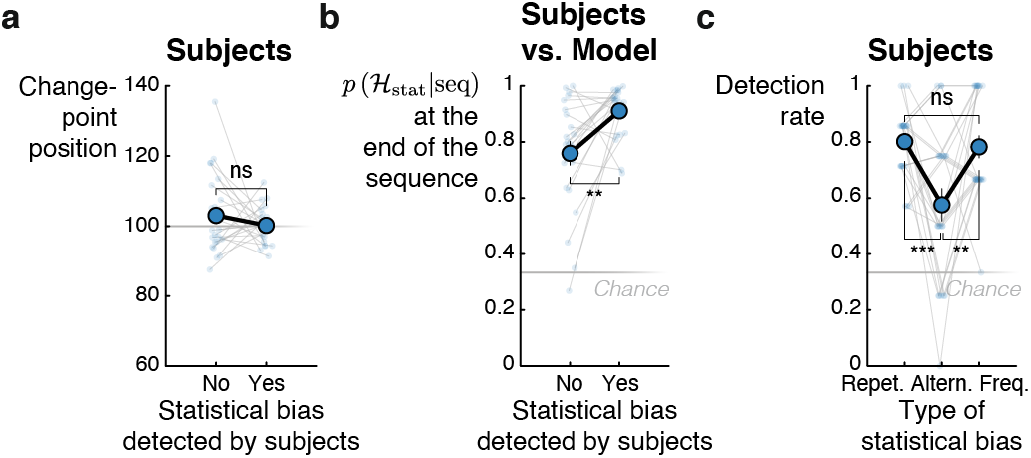
Normative causes of undetected statistical biases. **a**, *No latter change-point occurrence*. Sequences with undetected statistical biases were not characterized by a latter occurence of the change-point. **b**, *Weaker evidence*. Sequences with undetected statistical biases were characterized by a lower posterior probability of the statistical bias hypothesis estimated by the model at the end of the sequence. **c**, *Less frequent detection of alternation biases*. For both the subjects and the model, alternation biases are more often missed than repetition- and frequency-biases (after controlling for the strength of statistical bias, by design). This is a signature of transition probability learning. Stars denote significance: *** *p* < 0.005, ** *p* < 0.01, * *p* < 0.05; ns stands for non-significant.

**Extended Data Fig. 4.**
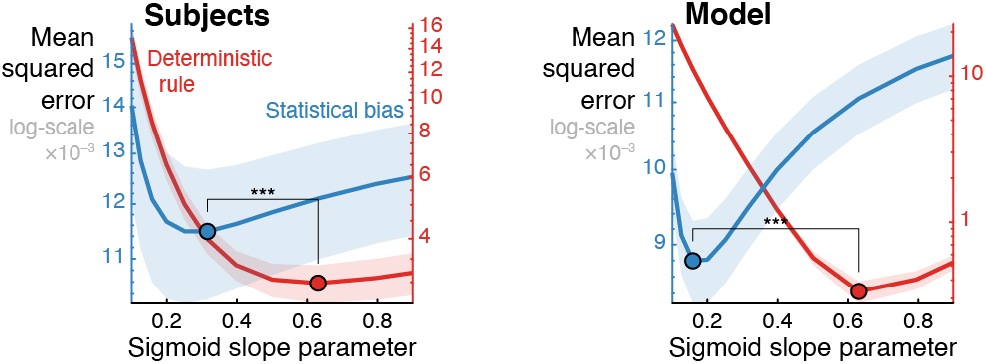
Different detection slope for statistical biases and deterministic rules. The distribution of mean squared error (MSE) is plotted as a function of a grid of values of the sigmoid slope parameter (ater minimizing the MSE over the other parameters of the sigmoid functions). Analyses were restricted to non-random sequences that were correctly identified by subjects and shaded areas correspond to the standard error of the mean computed over subjects. Stars denote significance: *** *p* < 0.005, ** *p* < 0.01, * *p* < 0.05.

**Extended Data Fig. 5.**
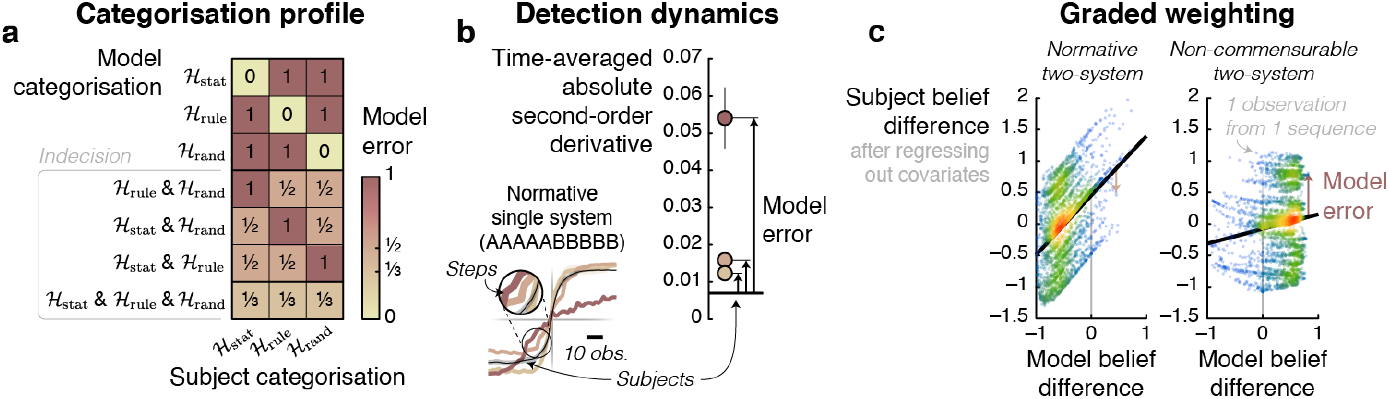
Error metrics. **a**, *Error measured on categorisation profile*. Model error is measured by comparing the model’s and subject’s categorisation of sequences. Subjects report the generative process using post-sequence questions. Models identify the most likely generative process based on the posterior probability of each hypothesis at the end of the sequence. When models were undecided (i.e. when two hypotheses have probability ½± 0.001, or three hypotheses have probability ⅓ ± 0.001; see bottom part of the matrix), the resulting error is divided among the equally likely hypotheses. **b**, *Error measured on detection dynamics of deterministic rule*. Model error is measured by comparing the model’s and the subject’s detection dynamics of deterministic rules. A metric reflecting the smoothness of the dynamics is used: the absolute second-order derivative of the posterior probability of the deterministic rule hypothesis averaged in a period ranging from the change-point position to the end of the sequence. Each sequence (with a deterministic rule) is thus characterized by such a metric. **c**, *Error measured on graded weighing of non-random hypotheses in random sequences*. Model error is the residual error of the linear regression relating subject’s belief difference in random sequences to the model’s, restrained to ≥ 0 regression coefficients to prevent flips of data points between models and subject. The effect of observation number and together with its interaction with belief difference (*Fig. 7c*) is removed prior to this analysis (using a linear regression).

**Extended Data Fig. 6.**
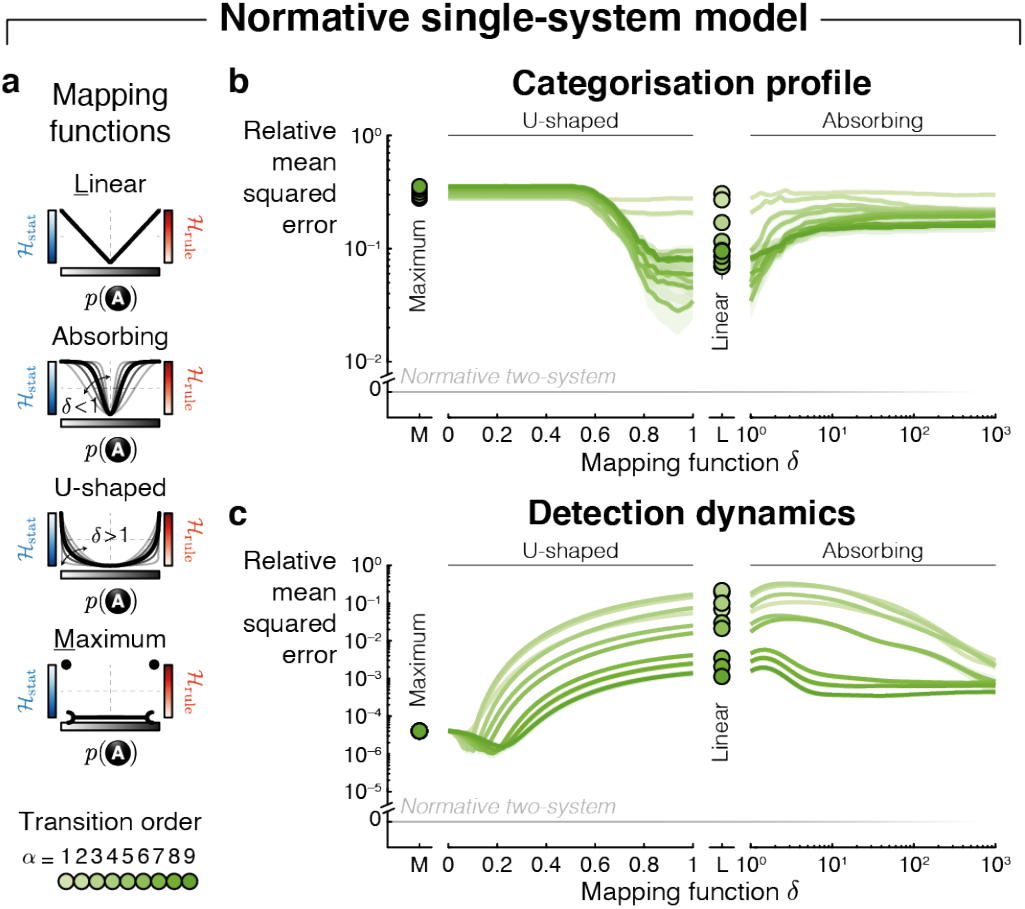
*Normative single-system model* arbitrating between non-random hypotheses using non-linear functions of predictability strength. **a**, *Non-linear functions of predictability strength*. In the version of the *normative single-system model* in which *H*_stat_ and *H*_rule_ monitor the same transition order (which we varied) using the same prior beliefs regarding predictability (flat prior), we explored several variants. Because, in this case, the two non-random hypotheses are strictly identical, the strength of predictability is used to arbitrate among non-random hypotheses: the further away *p*(*y_k_*|*y*_1:k−1_), where *y* is the sequence and *k* the current observation, from ½, the more likely *H*_rule_. In the main text, we report results for an estimation of posterior probabilities estimated as a linear function of predictability. Here, we explored non-linear relationships: maximum, U-shaped, and absorbing functions. **b**/**c**, *Error of alternative models with respect to subjects’ categorisation profile/detection dynamics of deterministic rules*. Same as in *Fig. 8*.

**Extended Data Fig. 7.**
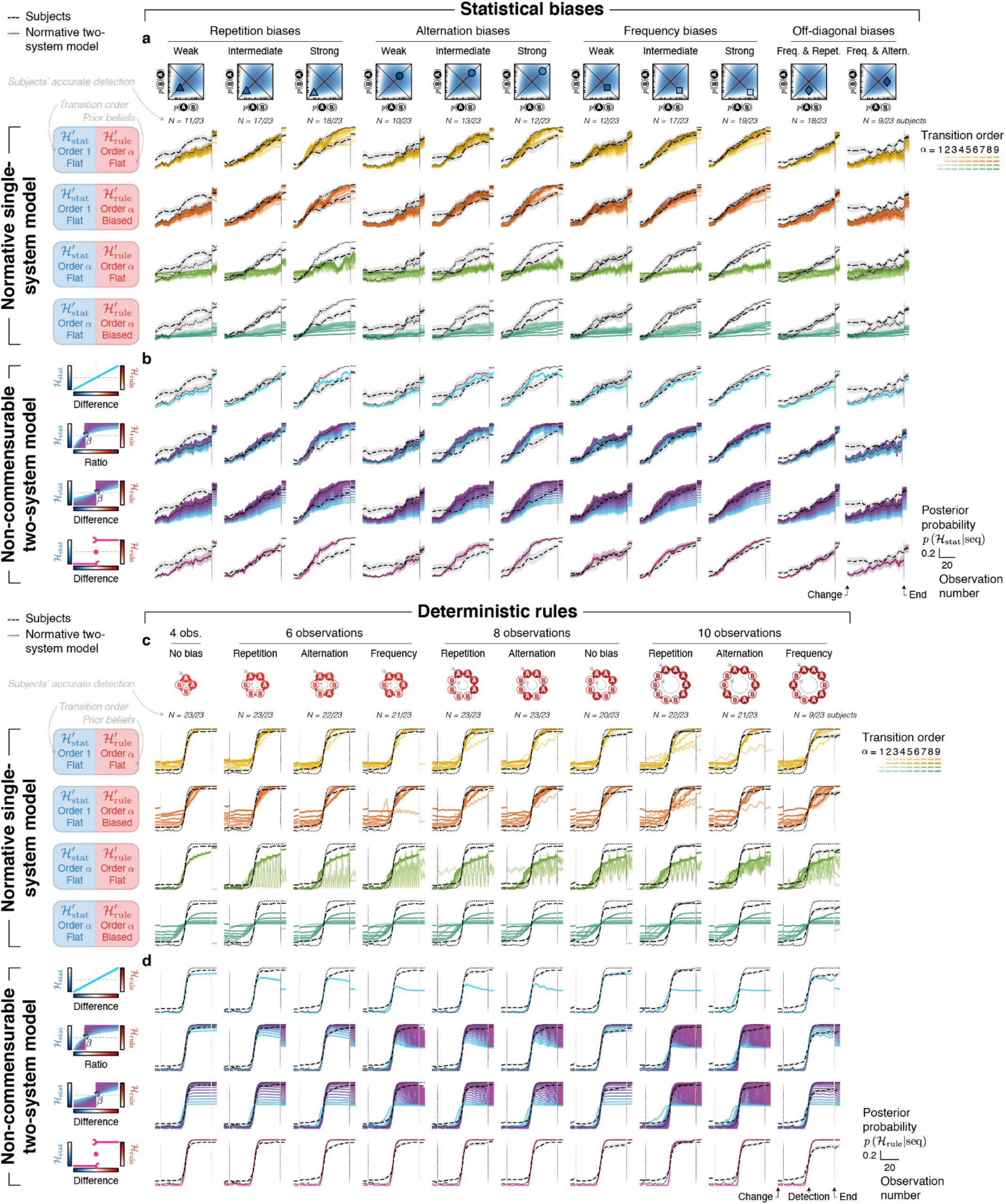
Detection dynamics by subjects and models. **a**/**b**, *Detection dynamics of statistical biases from the normative single-system model/non-commensurable two-system model*. The posterior probability of the statistical bias hypothesis is aligned on the change-point position. Detection dynamics is plotted for each deterministic rule (columns), for each version of the models (rows), and across a range of parameter values (color-coded lines). Detection dynamics from the *normative two-system model* and the subjects are overlaid on all plots as dotted and dashed lines respectively. **c**/**d**, *Detection dynamics of deterministic rules from the normative single-system model/non-commensurable two-system model*. The posterior probability of the deterministic rule hypothesis is aligned on detection-point. Same convention as in **a**/**b**.

**Extended Data Fig. 8.**
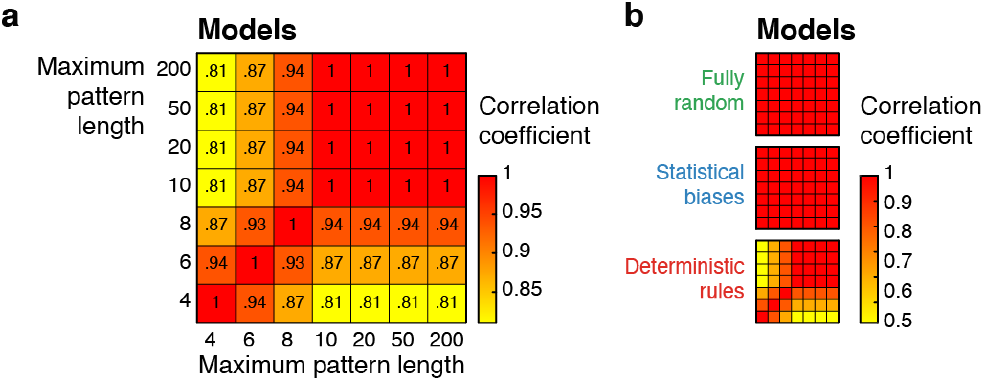
Similarity of model inference for different maximum pattern lengths. **a**, *Correlation across all types of sequences*. Correlation between posterior probabilities (after conversion to cartesian coordinates to ensure an appropriate number of degrees of freedom) from different versions of the *normative two-system model* considering different maximum lengths for pattern detection. Versions of the model which use a maximum pattern length larger than the longest patterns (i.e. 10 observations) used in the experiment make very similar inferences. **b**, *Correlation across each type of sequence*. Same as in **a** but separately for each type of sequence. The difference in inference between versions of the model using small maximum pattern lengths arise solely for sequences entailing deterministic rules, thereby suggesting that the difference solely originates from longest patterns remaining undetected.

## Supplementary information

### Supplementary note 1 — Subjects’ inference quantitatively deviate from optimality in several ways

Hereafter, the model we refer to is the *normative two-system model*.

Even though many aspects of human behaviour are well accounted for by the *normative two-system model* (see *Supplementary Fig. 1* for a summary), this does not mean that human behaviour is fully optimal. Instead, we observed several quantitative deviations from optimality.

**Supplementary Fig. 1.**
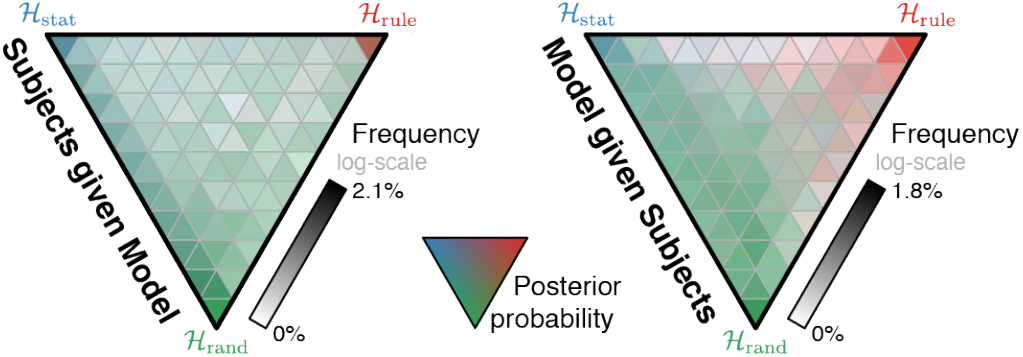
Overall agreement between subjects and the model. The agreement map on the left panel depicts what was the hypothesis posterior probability reported by subjects (which is color-coded according to the triangular colormap) as a function of the corresponding hypothesis posterior probability from the model (averaged in 10 × 10 × 10 equally spaced bins). Transparency reflects the normalized log-number of observations in each bin. Right panel shows the same thing but for the model conditionally upon what the subjects reported. These maps are obtained by concatenating data from all subjects and all sequences together. The maps show strong agreement between subjects and the model both for extreme (corresponding colours at each summit) and intermediate (progressively changing colours in between pairs of summits) posterior probabilities.

#### Integration delay

Because of the presentation rate (one sound every 300 milliseconds), it seems a priori difficult for subjects to update immediately their reported likelihoods, thereby inducing a delay between the presentation of the observation and its effect on the reports. We estimated correlations between posterior probabilities reported by human subjects vs. the model (after conversion to cartesian coordinates to ensure an appropriate number of degrees of freedom), after shifting in time one with respect to the other, in order to estimate the most likely integration delay characterizing each subject. Integration delay was found to be significantly larger than zero (delay = 5.00 observations ≈ 1.5 second, CI = [3.49, 6.51], *d*_Cohen_ = 1.43, *t*_22_ = 6.87, *p* = 6.70 × 10^−7^; *Supplementary Fig. 2a*). Interestingly, we also found a positive correlation between the integration delay and the correlation with the model when taking into account the delay (*r*_21_ = 0.74, *p* = 4.96 × 10^−5^; *Supplementary Fig. 2b*), suggesting that more cautious subjects reported more accurate values.

**Supplementary Fig. 2.**
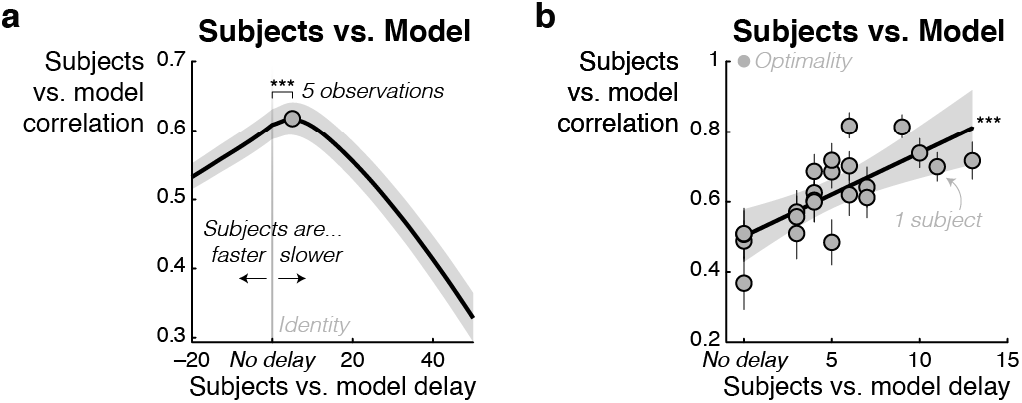
Delay between subjects’ and model’s reports. **a**, *Response delay*. Coefficients from a cross-correlation between all hypothesis posterior probabilities from subjects and the model (after conversion to cartesian coordinates to ensure an appropriate number of degrees of freedom) after shifting in time one with respect to the other. The average delay corresponds to ~5 observations. Error bars correspond to the standard error of the mean computed over subjects. **b**, *Inter-subject correlation between estimated delay and correlation strength with the model*. Subjects with longer integration delay reported more accurate values. Error bars correspond to the standard error of the mean computed for each subject over sequences and the shaded area to 95% confidence interval of the regression coefficients. Stars denote significance: *** *p* < 0.005, ** *p* < 0.01, * *p* < 0.05.

#### Probability weighting

The literature on probabilistic inference frequently reports that humans overestimate small probabilities and underestimate large probabilities (*Supplementary Fig. 3a*), a phenomenon termed probability weighting and which is commonly described by the following function which maps objective (*p*) to subjective (*p*^*^) probabilities

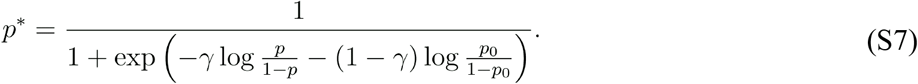

To assess and quantify the distortion (relative to optimality) of subjects’ reported posterior probability in the different hypotheses, we estimated the mean squared error (MSE, relative to a linear mapping) between posterior probabilities reported by human subjects vs. the model (using cartesian coordinates and not barycentric ones to use an appropriate number of degrees of freedom). More precisely, we used a grid from 0.01 and 1.99 for the slope parameter (*γ*) and from 0.01 to 0.99 for the offset parameter (*p*_0_), with 0.01 steps in both cases, and found the set of parameters inducing the smallest MSE. We found the classic under- /overestimation effect in our data: the slope parameter was found to be significantly smaller than 1 (difference of *γ* slope parameter to 1 = −0.55, CI = [−0.67, −0.43], *d*_Cohen_ = −2.01, *t*_22_ = −9.65, *p* = 2.29 × 10^−9^; *Supplementary Fig. 3b*) and induced a MSE significantly smaller that an unbiased linear mapping (difference in MSE = −0.078, CI = [−0.031, −0.015], *d*_Cohen_ = −1.21, *t*_22_ = −5.82, *p* = 7.38 × 10^−6^).

**Supplementary Fig. 3.**
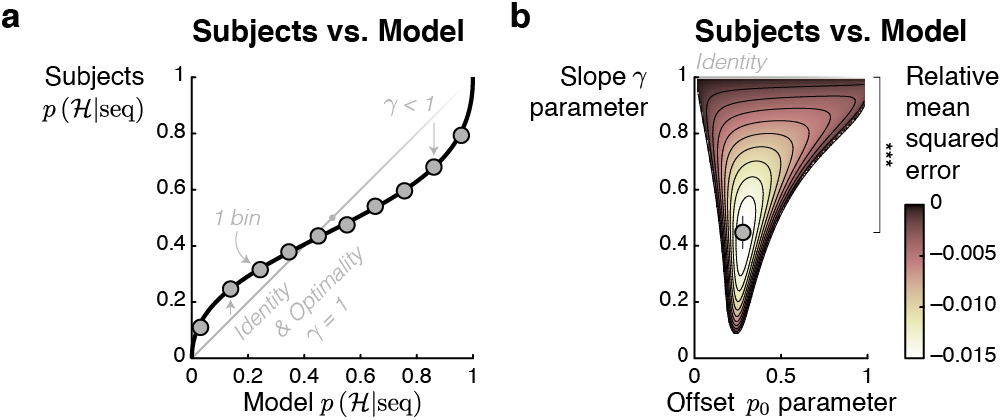
Non-linear mapping between subjects’ and model’s posterior probabilities. **a**, *Probability weighting*. All hypothesis posterior probabilities from the subjects are plotted against corresponding hypothesis posterior probabilities from the model (in 10 equally spaced bins). A nonlinear probability weighting function maps this relationship: small probabilities are overestimated by subjects and large ones are underestimated, leading to a *γ* slope parameter smaller than 1. Error bars correspond to the standard error of the mean computed over subjects. **b**, *Best parameter set*. Parameters of the probability weighting function (i.e. the slope *γ* and the offset *p*_0_) were varied along a grid of values. Mean squared error (i.e. MSE) between transformed probabilities (with the probability weighting function) from the model and corresponding probabilities from the subjects was computed (after conversion to cartesian coordinates to ensure an appropriate number of degrees of freedom). Here, the MSE is plotted relative to the error obtained with a linear, non-weighted, mapping (i.e. *γ* = 1). The grey dot represents the average best parameter set (i.e. which induces the smallest error). Parameter sets for which the linear mapping provides a better account (i.e. relative MSE > 0) have been masked. Stars denote significance: *** *p* < 0.005, ** *p* < 0.01, * *p* < 0.05.

### Supplementary note 2 — The explicit representation of change-point progressively discards past sequence observations from the inference

In past studies, we and others have emphasized the need to consider a memory leak when modeling human behaviour during sequence processing ^22,23,24,69,70^. It should be acknowledged that the models considered here do not rely on such a memory decay of past observations. Instead, the models explicitly represent the existence of a change-point. This means that the speed at which past observations are discarded from the inference depends upon the posterior beliefs about the position of the change-point: if it is likely that the change-point has occurred very recently then recent observations have more weight on the inference than remote ones. Marginalizing over possible change-point positions (with different posterior probabilities) progressively down-weights the more remote observations, similarly to the memory leak described in previous studies, but with the difference that such down-weighting is adaptive in change-point models vs. fixed in models with memory leak.

In order to quantify the strength of such a leak in the model, we computed the equivalent learning rate:

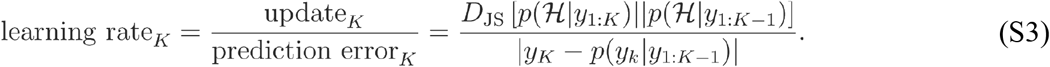

On the one hand, the successive update between posterior distribution over hypotheses is computed using the Jensen-Shannon divergence (i.e. *D*_JS_; Lin, 1991), which corresponds to

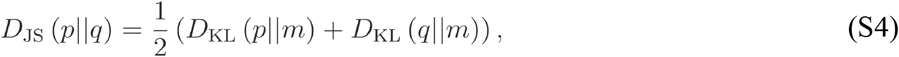

with 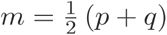, and where the Kullback-Leibler divergence (i.e. *D*_KL_) is defined as

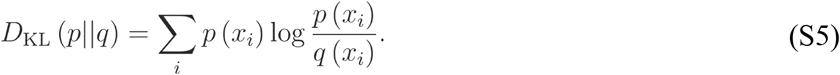

Compared to *D*_KL_, *D*_JS_ has the advantage to be symmetric, that is *D*_JS_(*p*∥*q*) = *D*_JS_(*q*∥*p*); note however that similar conclusions are reached with *D*_KL_. On the other hand, the model prediction (necessary to compute the prediction error term) is

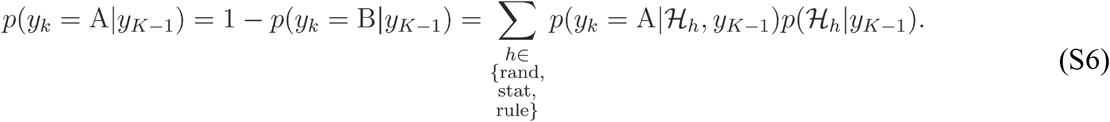

The larger the (equivalent) learning rate, the faster previous observations are down-weighted (*Supplementary Fig. 4a*). Inspecting the distribution of equivalent learning rates in all sequences reveals that the model uses a range of equivalent learning rates (*Supplementary Fig. 4b*). In particular, the equivalent learning rate increases following the change-point. This increase furthermore reflects the slope of the detection: larger increase of equivalent learning rate for detection rules than statistical biases (*Supplementary Fig. 4c*).

**Supplementary Fig. 4.**
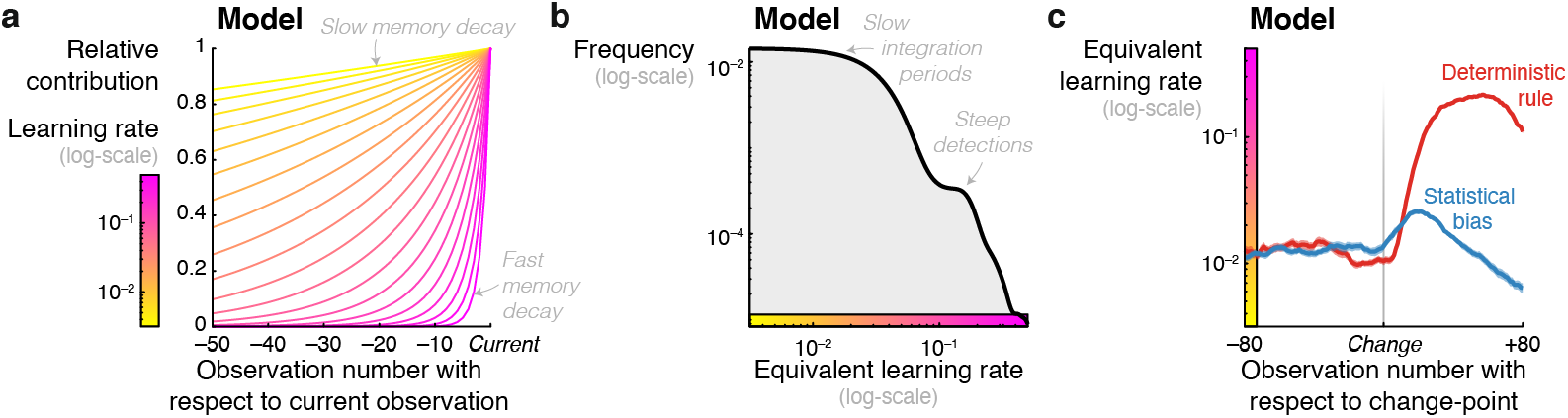
A variety of learning rates. **a,** *Learning rate determines the strength of memory decay*. Stronger learning rates lead to faster decrease in the relative contribution of past observations, that is a stronger memory decay. **b,** *Distribution of equivalent learning rate*. Model’s equivalent learning rates are estimated from the prediction error and the amount of update evoked by each sequence observation. The distribution of equivalent learning rates across all observations from all sequences are estimated using a gaussian kernel density with a bandwidth of 0.025. The distribution reveals a variety of equivalent learning rates used by the model, thereby reflecting different forgetting regimes. **c,** *Increase of equivalent learning rate following change-point*. Equivalent learning rate from the model were locked on change-points and averaged over sequences separately for random-to-statistics and random-to-rule sequences, with a temporal smoothing (moving mean) of 20 observations. In **c**, analyses were restricted to non-random sequences that were correctly identified by subjects and for which detection-points were found for both the subjects and the model. In **b** and **c**, the shaded area corresponds to the standard error of the mean computed over subjects.

